# “Peptidergic modulation of motor neuron output via CART signaling at C bouton synapses”

**DOI:** 10.1101/2022.11.05.515234

**Authors:** Panagiotis E. Eleftheriadis, Konstantinos Pothakos, Simon A. Sharples, Panagiota E. Apostolou, Maria Mina, Efstathia Tetringa, Gareth B. Miles, Laskaro Zagoraiou

## Abstract

The intensity of muscle contraction, and therefore movement vigour, needs to be adaptable to enable complex motor behaviors. This can be achieved by adjusting the properties of motor neurons, which form the final common pathway for all motor output from the central nervous system. Here we identify novel roles for a neuropeptide, Cocaine and Amphetamine Regulated Transcript (CART), in the control of movement vigour. We reveal distinct, but parallel mechanisms by which CART and acetylcholine, both released at C bouton synapses on motor neurons, selectively amplify the output of subtypes of motor neurons that are recruited during intense movement. We find that mice with broad genetic deletion of CART or selective elimination of acetylcholine from C boutons exhibit deficits in behavioral tasks that require higher levels of motor output. Overall, these data uncover novel spinal modulatory mechanisms that control movement vigour to support movements that require a high degree of muscle force.

## Introduction

The generation of complex motor behavior is critically dependent on the ability to precisely time the sequence of muscle activation and to regulate the amount of force produced by muscles. Whilst the timing of muscle activation is largely encoded by integrated populations of spinal interneurons ^1,2^, the strength of muscle activation, and subsequent movement vigour, is determined by motor neuron output. Importantly, motor neuron output is constantly adjusted to suit the task-dependent demands of diverse motor behaviors. These adjustments are often achieved by regulating cellular properties that determine motor neuron recruitment and firing rates ^3–5^. A range of neuromodulatory pathways, originating from higher centers and local interneurons, are responsible for this regulation of motor neuron properties. One of the most prominent spinally-derived neuromodulatory pathways for the regulation of motor output is mediated by spinal V0c interneurons and their C bouton synapses at motor neurons ^6,7^.

The broad V0 population of spinal interneurons, which are identified based on the expression of the Developing Brain Homeobox 1 transcription factor (Dbx1), were one of the first developmentally-defined populations shown to provide direct input to motor neurons ^8^. The V0 population, is composed of dorsal and ventral subpopulations ^9,10^. The ventral subpopulation includes a small subset that can be identified based on the expression of the Paired-like homeodomain transcription factor 2 (Pitx2), which is further subdivided into cholinergic (V0c) and glutamatergic (V0g) subsets ^7^. Although V0c interneurons comprise only 2.5% of the cardinal V0 population, they represent the sole source of large cholinergic C bouton synapses at the soma and proximal dendrites of motor neurons. V0c interneurons, and their C boutons, support the key function of increasing motor output during tasks of increased motor demand ^7^. Although significant knowledge exists regarding the pre- and post-synaptic machinery of the C bouton synapse, the mechanisms by which this system adjusts motor output remain to be fully resolved. The traditional view is that the release of acetylcholine from C boutons increases motor neuron firing rates through the activation of postsynaptic M2 muscarinic receptors leading to downstream effects on a range of ion channels including calcium-activated potassium channels and Kv2.1 channels ^7,11^.

Here, we reveal novel mechanisms by which V0c interneurons, and their C bouton synapses, regulate motor neuron activity. We show that a poorly understood signalling neuropeptide, the Cocaine and Amphetamine Regulated Transcript (CART), is expressed in V0c interneurons at all levels of the spinal cord and in most C boutons. Functional analysis reveals distinct, yet parallel mechanisms for CART and acetylcholine in the amplification of the output of a subset of motor neurons, fast motor neurons, that are typically only engaged during tasks that require high levels of motor output ^12^. We found that CART facilitates the recruitment of fast but not slow motor neurons, whereas cholinergic pathways increase firing rates of fast motor neurons. Intriguingly, neither CART nor acetylcholine are critical for the formation or maintenance of C bouton synapse structure, but they are important for the generation of motor behaviors that require high levels of motor output. Together, these findings support that CART represents a novel parallel signalling mechanism at this important neuromodulatory synapse. Given that cholinergic modulation of output neurons is a common feature of neural networks throughout the CNS^13–15^ our results are likely to be applicable to a range of networks and behaviors.

## Results

### Cocaine and Amphetamine Regulated Transcript is enriched in a subset of Pitx2 spinal interneurons

With the aim to identify markers to distinguish homogeneous, functionally-defined subsets of interneurons involved in motor control and locomotion we have previously performed a microarray screen ^7^ comparing the mRNA expression in the ventral and dorsal parts of the lumbar spinal cord of mice at postnatal day 8 (P8; Fig. 1a). We focused on genes that are preferentially expressed in the ventral, lumbar spinal cord where locomotor circuits reside ^16^. This screen resulted in the identification of 82 genes that were expressed more than three-fold higher in the ventral compared to the dorsal horn ^7^ two of which, Pitx2 and Short-stature homeobox 2 (Shox2), have been previously described and are expressed exclusively in subsets of spinal interneurons that play key roles in the control of motor neuron excitability and locomotor rhythm generation respectively ^7,17^. This study focuses on one of the genes highlighted by our screen, the *Cart* gene, one of the most enriched transcripts in the ventral spinal cord, which encodes a signaling peptide called the Cocaine and Amphetamine Regulated Transcript (CART) that following in situ hybridization was revealed to be expressed in a small interneuron subset in the intermediate zone of the spinal cord. CART was found to be expressed in a 3.4:1 ratio in the ventral compared to dorsal horn. *Cart* mRNA expression in P8 mice was detected in neurons located near the central canal, in the motor neuron area (Laminae VIII and IX), and the dorsal horn of the cervical (Fig. 1b), thoracic (Fig. 1c), and lumbar (Fig. 1d, e) segments of the spinal cord. In addition, *Cart* expressing neurons were also detected in the intermediate zone of the thoracic spinal cord, representing sympathetic preganglionic neurons (SPNs) (Supplementary Fig. 1a), consistent with previous reports ^18–20^. A similar expression profile was observed in the adult thoracic spinal cord (P25), where *Cart* mRNA was observed near the central canal, in the motor neuron-area, in putative SPNs, and in the dorsal horn (Fig. 1f).

**Fig. 1:**
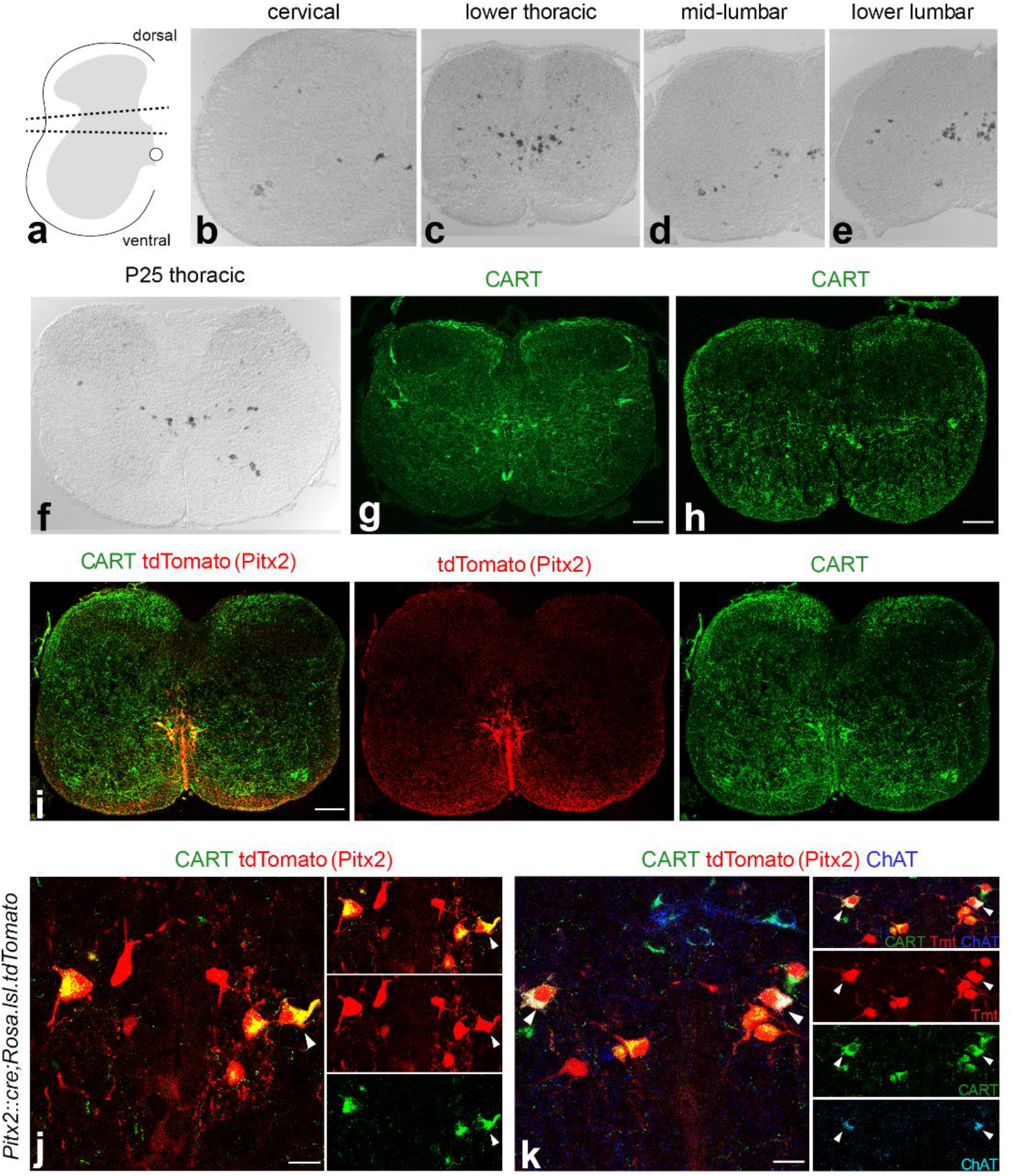
CART and Pitx2 expression in the spinal cord partially overlap. **(a)** Schematic representation of the tissue dissected for the ventral vs. dorsal horn microarray screen. Tissue between dotted lines was discarded. (**b-e**) *Cart* mRNA detection in all levels of P8 mouse spinal cord as revealed by *in situ* hybridization. (**f**) *Cart* mRNA detection in P25 spinal cord. The presence of *Cart* mRNA is mainly concentrated in the peri-central canal region as well as, in some motor neuron somata. (**g, h**) CART neuropeptide detection in spinal cord cross sections of P8 (**g**) and P0 mice (**h**). (**i**) Partial co-detection (yellow) of CART (green) and tdTomato (red) in the Pitx2 expressing neurons from spinal cords of *Pitx2::Cre*;*Rosa*.*lsl*.*tdTomato* mice (P0). (**j**) High magnification of the peri central canal region, cropped neurons presented in separate panels [merge, tdTomato (red), CART (green)]. Some tdTomato+ neurons co-express CART (tdTomato+/CART+ neurons), only one is indicated by a white arrowhead for simplicity; tdTomato+/CART-neurons are also observed (P0). (**k**) V0c cholinergic neurons co-express CART (tdTomato+/ChAT+/CART+, red/blue/green respectively, white arrowheads). Cropped neurons presented in separate panels [merge, tdTomato (red), CART (green), ChAT (Cyan)]. Non-cholinergic (V0g) Pitx2 neurons can be subdivided to CART expressing (tdTomato+/CART+/ChAT-, red and green) and non-expressing (tdTomato+/CART-/ChAT-, red only) (P0). Scale bars = 100 μm (**g-i**), 20μm (**j, k**).

We next performed immunohistochemistry experiments to examine the distribution of the CART peptide within the spinal cord. Consistent with in situ hybridization, CART-labelled somata were identified near the central canal, in the motor neuron area, in regions where SPNs reside, and in the dorsal horn of the P8 upper lumbar spinal cord (Fig. 1g). CART expressing neurons, as revealed by immunohistochemistry, recapitulated the in situ hybridization data in all spinal segments. Interestingly, we not only detected CART immunoreactivity in somata, but also in fibers in the superficial dorsal horn, Laminae VII-X, and through the anterior white commissure. A similar distribution of CART expression was also observed in spinal cords obtained from mice at P0 suggesting that CART expression is also present at birth, albeit with a lower intensity in the superficial dorsal horn (Fig. 1h).

We next set out to determine the identity of CART expressing neurons found near the central canal. Our attention was drawn to clusters of CART+ neurons located lateral to the central canal, that appear at all spinal levels, and were reminiscent of the V0cg (Pitx2+) subsets of interneurons. As shown in ^7^ the Pitx2 transcription factor defines two small interneuron subsets of the V0 population, the cholinergic V0c and the glutamatergic V0g, located in clusters close to the central canal and forming columns running throughout the rostrocaudal axis. The cholinergic V0c subset is known to be the sole source of cholinergic C bouton synapses on the somata and proximal dendrites of motor neurons. Therefore, should Pitx2 and CART expression co-localize a new marker and more importantly a potential neurotransmitter for the V0cg interneurons may be at hand.

To test this hypothesis, we crossed a *Pitx2::Cre* ^21^ mouse line with a *Rosa*.*lsl*.*tdTomato* ^22^ reporter line to express the fluorescent protein tdTomato in the tissues after excision of the LoxP-flanked stop cassette by Cre recombinase. CART was expressed in a subset of tdTomato-expressing somata (83%) (Fig. 1i,j), suggesting the CART peptide may only be expressed in a subset of the Pitx2+ interneurons. Surprisingly, 98% of V0c interneurons, co-labelled with Choline Acetyl Transferase (ChAT) and tdTomato, were CART+, specifically 97% in cervical, 98% in thoracic and 100% in upper lumbar levels (Fig. 1k). On the other hand, CART was only found in 65% of tdTomato+ ChAT-V0g interneurons; 65% in the cervical, 71% in thoracic and 61% in lumbar segments.

### Diverse expression profiles of CART within C bouton synapses

V0c interneurons give rise to large cholinergic synapses called C boutons that are located on the somata and proximal dendrites of alpha motor neurons and are important for facilitating output during tasks that require greater motor output ^7,11^. We therefore next examined CART expression at C boutons to determine if CART might be a novel signaling peptide at this synapse. Immunohistochemistry for CART demonstrated a high density of CART+ puncta in lamina VIII and IX. We also detected some CART+ motor neuron somata (ChAT+), as previously detected in the *in situ* hybridization experiments (Fig. 2a, Supplementary Fig. 2a, b). Focusing on lamina IX, the CART+ puncta were found to colocalize with ChAT+ terminals on motor neurons of P8 (Fig. 2b) and P25 mice (Fig. 2c), suggesting that CART may serve as a second neurotransmitter at C bouton synapses (Fig. 2d).

**Fig. 2:**
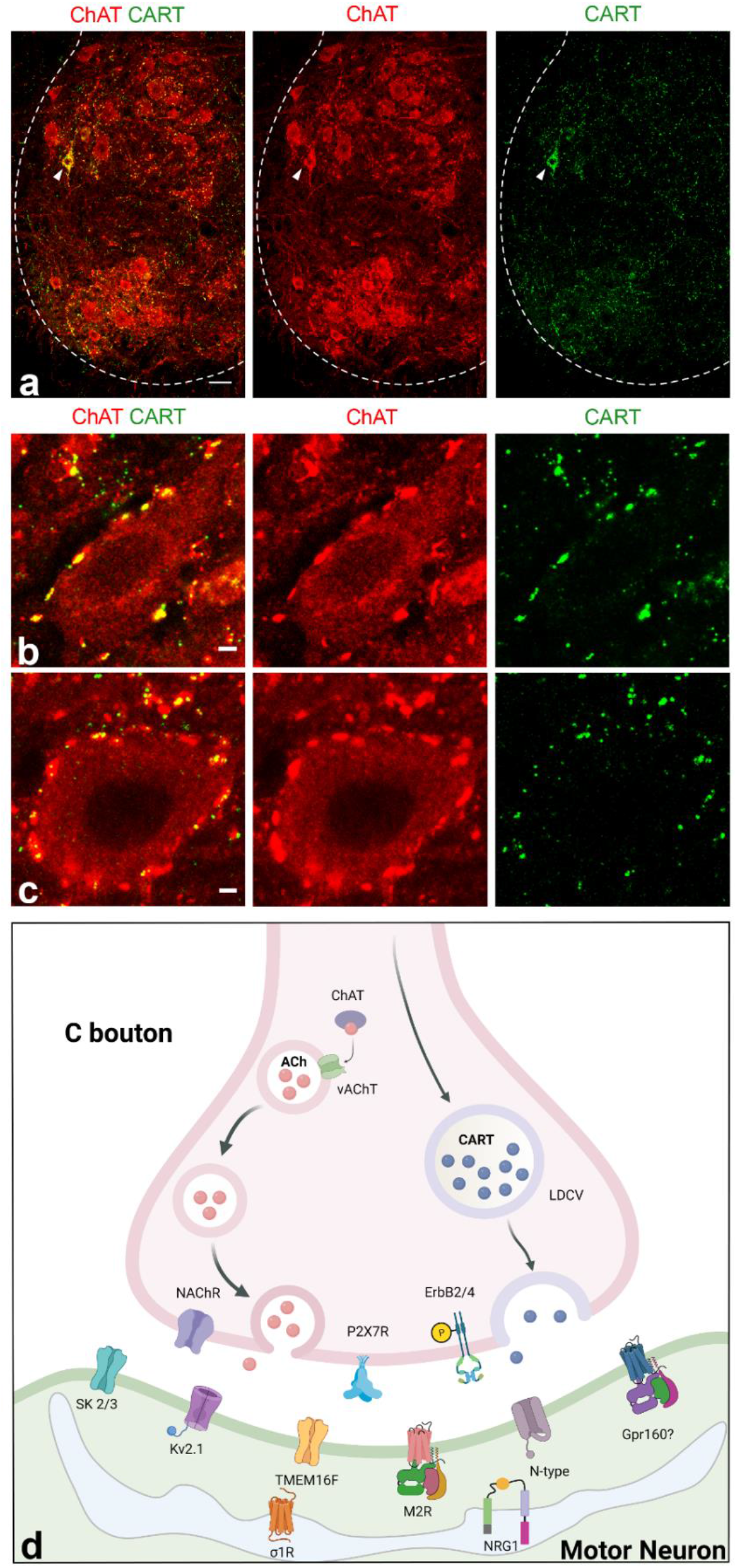
CART detection in C bouton synapses and in a subset of spinal motor neurons. **(a)** Immunohistochemistry in P25 mice with antibodies against ChAT (red) and CART (green) reveals the presence of CART puncta in the motor neuron area, specifically on motor neuron membranes. CART immunoreactivity is also observed in few motor neuron somata (white arrowhead). Scale bar=20 μm. (**b**) Co-localization of ChAT (red) and CART (green) in the cholinergic C bouton synapses on motor neurons of P8 mice. (**c**) C bouton synapses of P25 mice also contain CART. Scale bars = 2μm (**b**,**c**). (**d**) Schematic representation of the C bouton synapse (Created with Biorender). In the presynaptic area, the C boutons contain ChAT, vAChT, and CART as well as the presynaptic receptors P2X_7_R ErbB2/4 and NAChR. Post-synaptically, the M2 muscarinic receptors are present along with the SK2/3, N type, Kv2.1 and TMEM16F channel. The Gpr160 receptor is speculated to be located in the post-synaptic terminal as recent work suggests it is a putative CART receptor. In the subsurface cisterns sigma 1 receptor is also present along with NRG1.

Colocalization of CART and ChAT in C bouton synapses (Fig. 3a), and in some motor neuron somata (Fig. 3b) was further supported by additional experiments in spinal cord sections from *Pitx2::Cre;Rosa*.*lsl*.*tdTomato* mice, where C bouton synapses were visualized with tdTomato. Using the IMARIS Image Analysis software (Oxford Instruments) on deconvolved confocal images, two different 3D surfaces were created that outline the volumes of interest (tdTomato: red surface, CART: green surface) revealing the partial or complete enclosure of the CART surfaces in the synapse (Fig. 3c, Supplementary Fig. 3). Further observations of CART immunoreactivity patterns at C bouton synapses revealed a heterogenous distribution of CART within the presynaptic terminal. Synapses from spinal cords of *Pitx2::Cre;Rosa*.*lsl*.*tdTomato* mice and wild type mice (n=3 for each genotype, 400-700 synapses/mouse); labelled with anti-vesicular acetylcholine transporter (vAChT) and anti-CART antibodies were examined. Both approaches revealed four discrete distribution patterns of the CART neuropeptide in C bouton synapses (Fig. 3d). These included: i) “compact” synapses with an interior almost completely filled with the CART neuropeptide; ii) synapses with “puncta along the synapse” that had 3 or more CART+ puncta along the length of the synapse; iii) synapses with 1-3 puncta; and iv) also synapses with “peripheral puncta” that exhibited CART+ puncta at both ends of the synapse. Finally, we also observed cholinergic synapses that were devoid of the CART peptide. All CART distribution patterns were observed across postnatal development and through adulthood of mice (p10, p25, 8-9 weeks old, six months old, one year old; Fig. 3e), suggesting that this diversity is a general feature of CART at C bouton synapses.

**Fig. 3:**
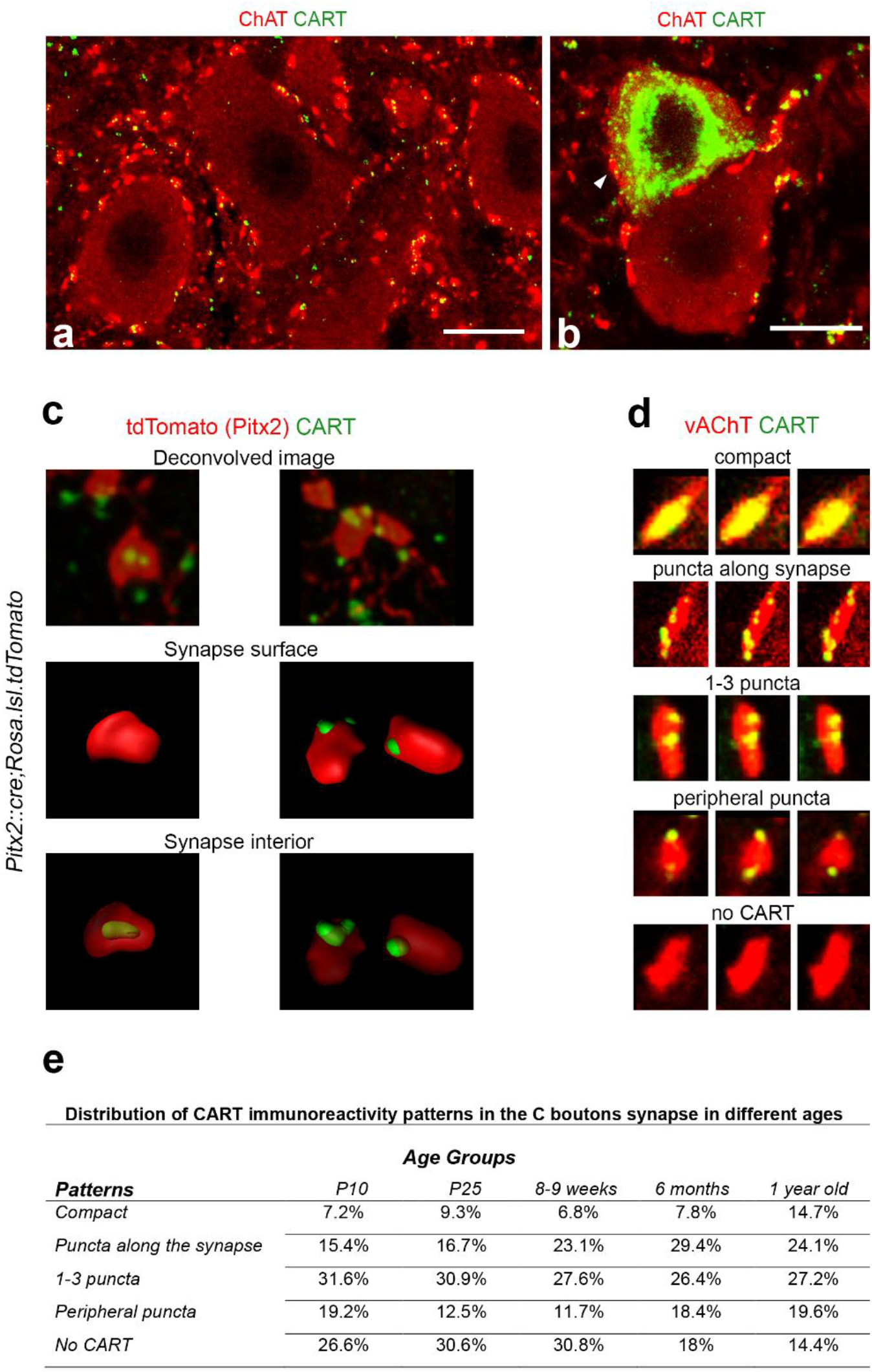
Patterns of CART immunoreactivity in C boutons. **(a)** ChAT+ (red) C bouton synapses on the somata of motor neurons of P25 mice contain CART (green). Scale bar = 20 μm. (**b**) CART (green) is also present in few motor neuron somata. CART+ motor neurons also receive C bouton synapses that contain CART (white arrow); (P25). Scale bar=20 μm. (**c**) Immunofluorescence with antibodies against CART (green) and dsRed (red) label tdTomato+ C boutons in spinal cord cross sections of *Pitx2:Cre*;*Rosa*.*lsl*.*tdTomato* P25 mice. Single C bouton synapses were 3D reconstructed using the IMARIS software. Upper panels: High magnification (deconvolved) confocal images, used for the 3D reconstruction, showing the CART neuropeptide (green) within the synapse (red). Middle panels (synapse surface): 3D reconstruction of the synapse surface (IMARIS). Lower panels (synapse interior): Rendering the synapse surface transparent reveals the distribution of CART inside the synapse. (**d**) The 5 different patterns of CART immunoreactivity in C boutons. Compact, puncta along synapse, 1-3 puncta, peripheral puncta and no CART. (**e**) Distribution of CART immunoreactivity patterns in the p10, p25, 8-9 weeks, 6 months old, 1 year old age groups as revealed by the analysis of n=3 mice per age group, where a mean of 544 synapses per motor neuron (1,208μm optical section thickness) were evaluated in lumbar segments (14um slices). The percentage of CART+ C bouton synapses differs depending on age.

### Parallel amplification of motor neuron output by CART and acetylcholine

We next sought to investigate whether CART provides an additional, physiologically relevant signalling mechanism at C bouton synapses. Given that C boutons are potent modulators of motor neuron excitability ^11,23^, we assessed whether CART could modulate the excitability of spinal motor neurons. We also compared any effects of CART with those of muscarine, which activates the predominant, known acetylcholine-based signaling mechanisms at C bouton synapses. We examined the effect of CART and muscarine on motor neuron recruitment and firing rate, which are two key mechanisms for the gradation of muscle force.

Previous work has reported higher densities of C boutons on fast compared to slow type motor neurons ^24^, in addition to differences in the postsynaptic structures located at C bouton synapses that contact different types of motor neurons ^25,26^. We therefore investigated whether CART or acetylcholine mediated modulation of the intrinsic properties that shape the recruitment or firing rate of motor neurons varied for fast versus slow type motor neurons from animals age P7-12. Motor neuron subtypes were identified based on their repetitive firing profiles ^27,28^ with fast motor neurons identified by a delayed firing and accelerating firing rate (Fig. 4a; n=21) and slow motor neurons identified by an immediate onset of repetitive firing and steady or adapting firing rates (Fig. 4b; n = 16). Consistent with these two populations representing fast and slow motor neuron types, and in line with previous work ^27,28^ delayed firing motor neurons had a lower input resistance (Fig. 4c; t(33)= 4.7, p = 4.3e-5) and a higher rheobase (Fig. 4c; t(34)=4.96, p = 2.1e-5), along with other differentiating intrinsic properties (summarized in Supplementary Fig. 4).

**Fig. 4:**
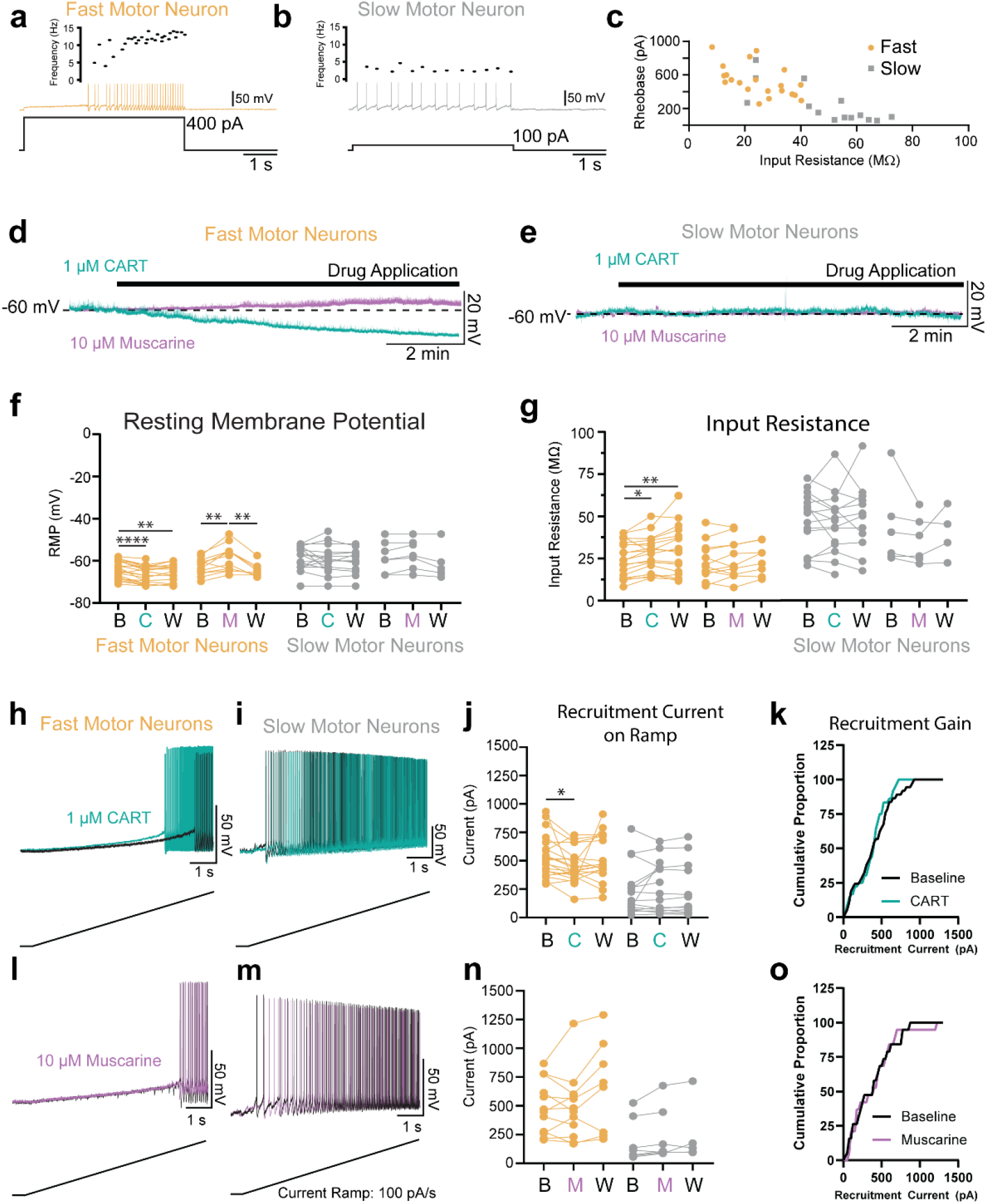
CART but not muscarine facilitates the recruitment of fast but not slow type motor neurons. (**a, b**) Representative trace of fast (**a**: orange) and slow (**b**: grey) motor neurons identified with whole cell patch clamp electrophysiology. (**c**) Scatter plot of input resistance and recruitment current (rheobase) demonstrate that fast motor neurons have a high rheobase and low input resistance, whereas slow firing motor neurons have relatively low rheobase and high input resistance. (**d, e**) Representative traces depicting the effects of CART (1μM; green) and muscarine (10 μM; purple) on the membrane potential of fast (**d**) and slow (**e**) motor neurons. (**f**) CART (C) and muscarine (M) produce opposing effects on the resting membrane potential (RMP) of fast (orange) motor neurons but do not alter the RMP of slow (grey) motor neurons. (**g**) CART (C) increases the input resistance of fast (orange) but not slow (grey) motor neurons. Muscarine (M) does not affect input resistance of fast or slow motor neurons. (**h, i**) Representative trace of the subthreshold voltage trajectory and onset of repetitive firing during a slow depolarizing current ramp (applied at 100 pA/s) in fast (**h**) and slow (**i**) motor neurons both before (black traces) and after application of 1 μM CART (green traces). (**j**) CART significantly reduces the recruitment current in fast (orange) but not slow motor neurons (grey). (**k**) Cumulative proportion histogram of motor neuron recruitment currents depicts an increase in the recruitment gain of the sample of motor neurons studied before (black) and after CART (green). (**l, m**) Representative trace of the subthreshold voltage trajectory and onset of repetitive firing in fast (**l**) and slow (**m**) motor neurons both before (black traces) and after application of 10 μM muscarine (purple traces). (**n**) Muscarine did not alter the recruitment current in fast (orange) or slow motor neurons (grey) and did not change the recruitment gain (**o**). All measurement were made at baseline (B), following drug application (CART (C), muscarine (M)), and following a wash period with regular aCSF (W). Asterisks denote significance (*p < 0.05, **p < 0.01, ***p < 0.001. ****p < 0.0001) from Holm-Sidak post-hoc following 2 factor (Repeated measures) ANOVA.

Having confirmed techniques to physiologically distinguish fast versus slow-type motor neurons, we next investigated the effects of CART (1 μM) or muscarine (10 μM) on the intrinsic properties of these motor neuron subtypes. Interestingly, both CART and muscarine selectively modulated several intrinsic properties in fast motor neurons only. Specifically, CART hyperpolarized the resting membrane potential (Fig. 4d, e, f; F (1.6,29) = 19.7, p = 2.2e-5), increased the input resistance (Fig. 4g; F (1.6, 30.6) = 7.6, p = 0.003), which translated to a reduction in the recruitment current (Fig. 4h, j; F (1.6, 30.6) = 3.5, p = 0.05), in fast but not slow type motor neurons (Fig. 4i, j). The decrease in recruitment current of the largest fast motor neurons led to an increase in the recruitment gain across the motor pool (Fig. 4k). Consistent with previous reports ^23,29^, we found that muscarine had a range of effects on motor neuron intrinsic properties. However, these effects were most robust in fast type motor neurons (n= 12), with no significant changes in the slow type motor neurons (n=7). In the fast motor neurons, muscarine depolarized the resting membrane potential (Fig. 4d, e, f; F (1.7,15.3) = 13.9, p = 5.4e-4), but did not change input resistance (Fig. 4g) or recruitment current of either fast or slow type motor neurons (Fig. 4l-o).

Both CART and muscarine also produced complementary modulation of fast but not slow motor neuron firing rates, assessed during slow depolarizing current ramps. CART increased firing rates at the lower end of the frequency current range (minimum firing rate: Fig. 5a, c; F (1.9,31.2) = 6.6, p = 0.005) and firing rate at 2x recruitment: Fig. 5d; F (1.8,29.7) = 3.4, p = 0.04), with no change in maximal firing rate (Fig. 5e; F (1.8,29.8) = 1.2, p = 0.3) in fast but not slow motor neurons (Fig. 5b, c-e). In contrast to CART, muscarine did not change the firing rate at recruitment (Fig. 5a, c; F (1.3,12.7) = 1.8, p = 0.2), but did increase the firing rate at 2x rheobase (Fig. 5d; F (1.4,13.2) = 9.2, p = 0.006) and the maximal firing rate (Fig. 5e; F (0.9,7.8) = 7.6, p = 0.03) in fast but not slow motor neurons (Fig. 5b, c-e). Both action potential rise time and half width were unaltered by CART or muscarine (Supplementary Tables 1, 2). CART and muscarine produced opposing control of the medium afterhyperpolarization (mAHP). CART increased the amplitude (Fig. 5f, h; F (1.2, 20.4) = 12.9, p = 0.001) and half width (Fig. 5k; F (1.6, 27.1) = 5.3, p = 0.02) of the mAHP of fast but not slow motor neurons (Fig. 5g, h, k). Muscarine reduced the mAHP amplitude (Fig. 5i, h; F (1.6,13.7) = 19.7, p = 2.3e-4) and half width (Fig. 5k; F (1.6,13.7 = 10.7, p = 0.002) in fast motor but not slow motor neurons (Fig. 5j, h, k).

**Fig. 5:**
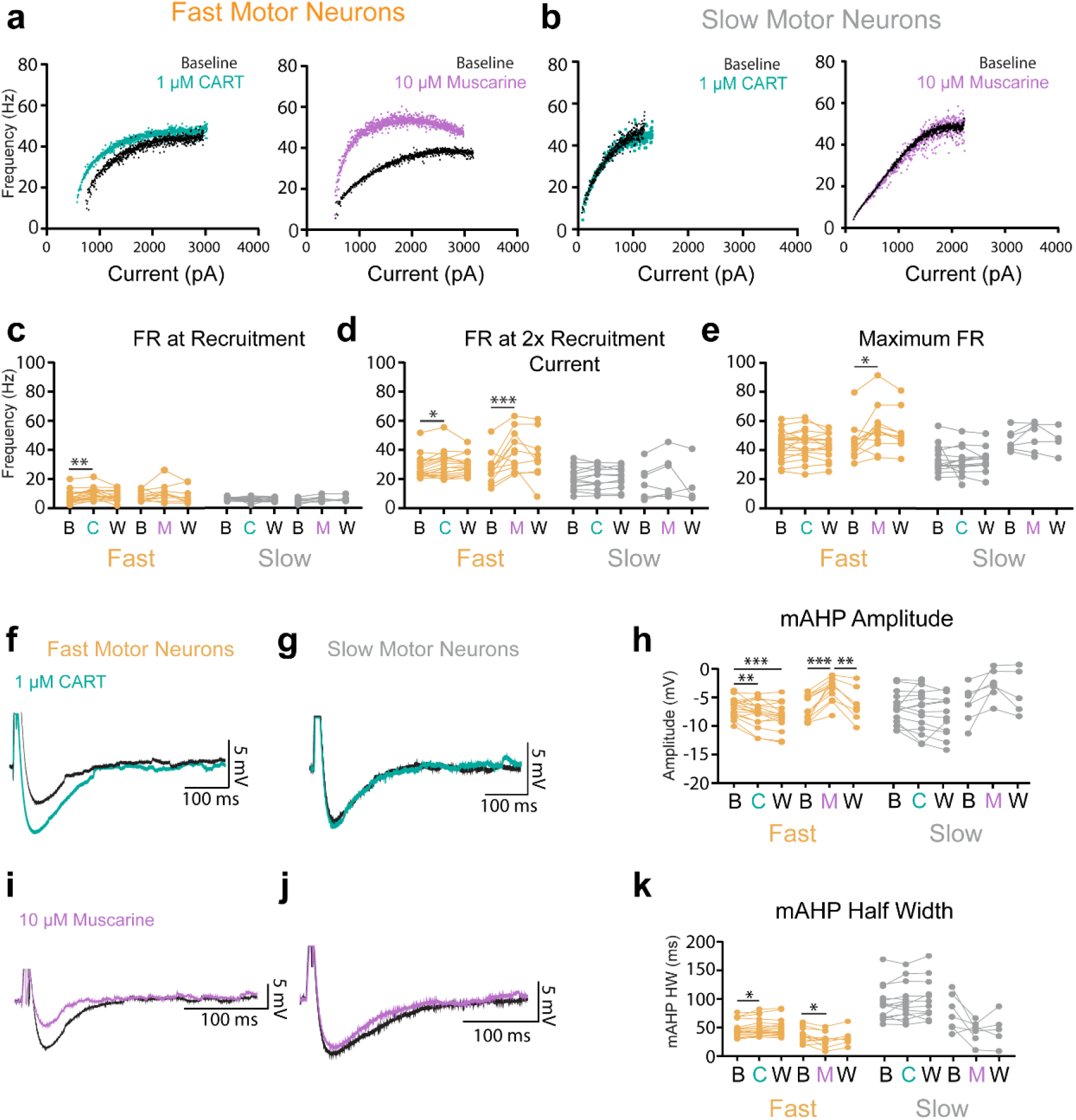
CART and muscarine modulate distinct components of frequency-current relationship and produce opposing control of mAHP in fast but not slow type motor neurons. **(a)**Representative frequency-current plots depicting changes in firing rate during a slow (100pA/s) depolarizing current ramp for fast motor neurons before (black) and after CART (green) or muscarine (purple). (**b**) Representative frequency-current plots depicting firing rates on current ramp for slow motor neurons before (black) and after CART (green) or muscarine (purple). (**c-e**) Effect of CART and muscarine on firing rates (FR) at recruitment, 2x recruitment current, and maximum. (**c**) CART significantly increases the firing rate of fast but not slow motor neurons at recruitment (minimum firing rate) and 2x recruitment current (**d**) but does not alter maximal firing rate (**e**), whereas muscarine does not alter firing rate for fast motor neurons at recruitment (**c**), but does increase firing rate at 2x recruitment current (**d**) and maximal firing rate (**e**). Neither CART nor muscarine alter firing rates of slow motor neurons. (**f, g**) Representative trace of the medium afterhyperpolarization (mAHP) in fast (**f**) and slow (**g**) motor neurons both before (black traces) and after application of 1 μM CART (green traces). (**h**) CART significantly increases whereas muscarine decreases the amplitude of the mAHP in fast motor neurons but do not alter the mAHP amplitude of slow motor neurons. (**i**,**j**) Representative trace of the mAHP in fast (**i**) and slow (**j**) motor neurons both before (black traces) and after application muscarine (purple traces). (k)CART (C) significantly increases whereas muscarine (M) decreases the half width (HW) of the mAHP in fast motor neurons but do not alter the mAHP HW of slow motor neurons. All measurements were made at baseline (B), following drug application (CART (C), muscarine (M)), and following a wash period with regular aCSF (W). Asterisks denote significance (*p < 0.05, **p < 0.01, ***p < 0.001. ****p < 0.0001) from Holm-Sidak post-hoc following 2 factor (Repeated measures) ANOVA.

Together, these data indicate that CART indeed affects motor neuron excitability. The two neurotransmitters, CART and acetylcholine, increase motor output through differential control of motor neuron subtypes via two key mechanisms that are important for the gradation of muscle force, with CART increasing recruitment and firing rates at the lower end of the input range and acetylcholine increasing firing rates throughout much of the input range without affecting recruitment or firing rate at recruitment.

### CART and acetylcholine are not essential for the formation and maintenance of C bouton structure

Having identified CART as a novel signaling molecule that is present alongside acetylcholine at C bouton synapses, we next set out to determine the relative importance of the two signaling pathways and what roles, if any, they may play in supporting the structural integrity of C bouton synapses. To address this, we studied C bouton synapses in P25 mice, identified with antibodies against vAChT and ChAT, when CART was globally knocked out ^30^ or ChAT conditionally deleted from V0c interneurons and their efferent C boutons (*Dbx1::Cre;ChAT*^*fl/fl*^),^31–33^(Fig. 6a-f). As expected, we detected no immunoreactivity for the CART peptide in CART^KO/KO^ mice (Fig. 6c). Strikingly, despite the absence of CART, C bouton synapses on motor neuron somata (ChAT+) remained well formed and appeared morphologically unaltered as confirmed by both ChAT and vAChT immunoreactivity (Fig. 6c). This result suggests that despite the complete absence of CART peptide, C bouton terminals remain structurally intact and develop in a manner consistent with control mice (Fig. 6a). The assessment of the conditional elimination of the ChAT gene from V0c interneurons yielded similar results (Fig. 6e). Despite the absence of the ChAT enzyme, CART peptide was still detected in C bouton synapses as revealed by vAChT labelling, suggesting that C bouton gross anatomy remains intact, and that CART peptide expression is unaltered and independent of the presence of acetylcholine (Fig. 6e; Schematic representations Fig. 6b, d, f).

**Fig. 6:**
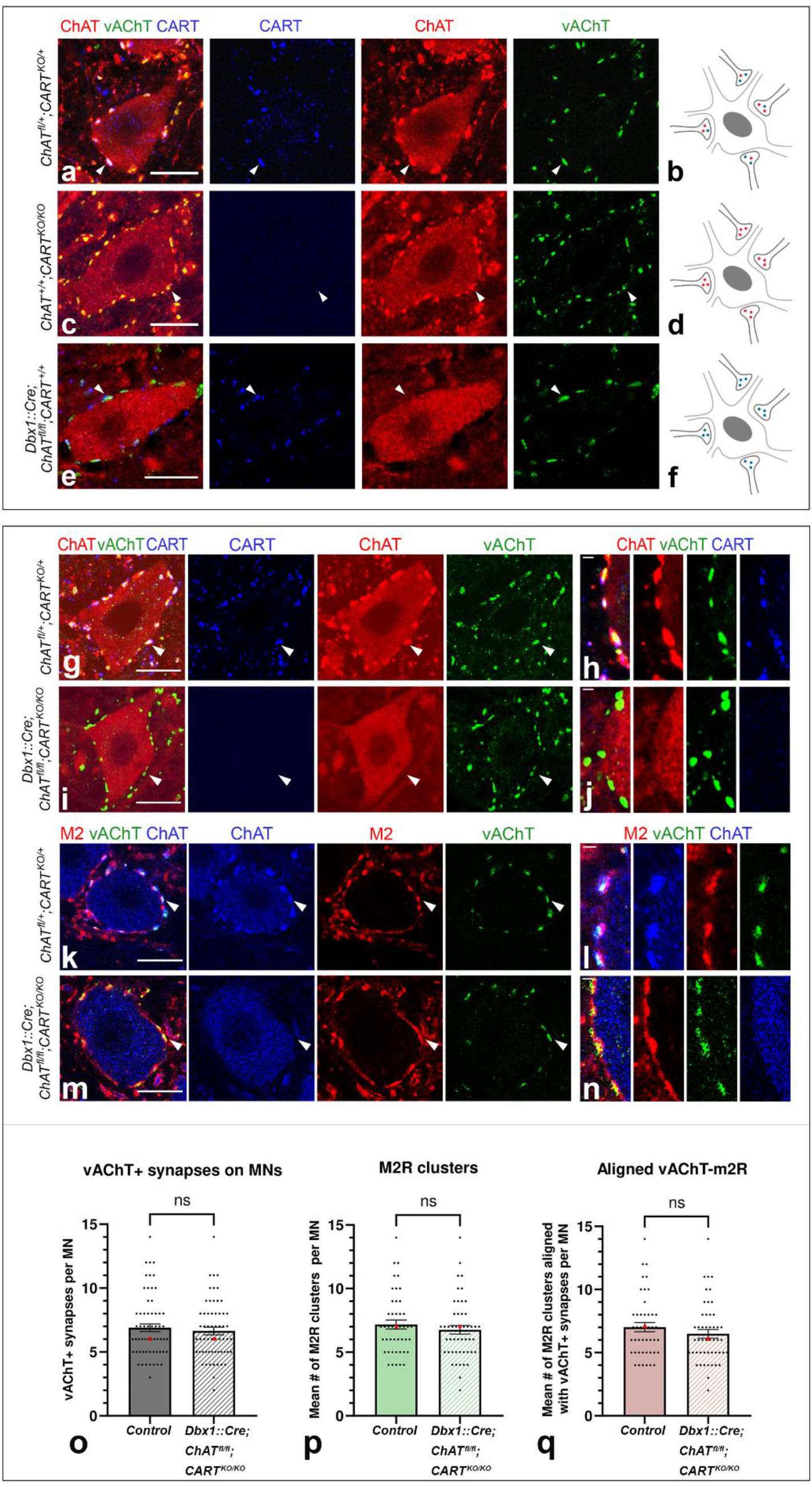
Gross anatomy of C boutons remains unchanged after CART and ChAT genetic elimination from the synapse. **(a)** C boutons on somata of motor neurons of wild type mice are marked by CART (blue) and the cholinergic markers ChAT (red) and vAChT (green). (**b**) Motor neuron sketch receiving C boutons containing acetylcholine (red) and CART (blue). (**c**) ChAT (red) and vAChT (green) expression are not disrupted in C boutons of *CART*^*KO/KO*^ mice. (**d**) Motor neuron sketch receiving C boutons containing only acetylcholine (red). (**e**) CART (blue) and vAChT (green) expression are not disrupted following genetic deletion of ChAT in V0c interneurons. (**f**) Motor neuron sketch receiving C boutons containing CART (blue) only. (**g**) CART (blue), ChAT (red) and vAChT (green) are present in C boutons of heterozygous *ChAT*^*fl/+*^*;CART*^*KO/+*^ mice. (**h**) High magnification image showing the colocalization of CART (blue), ChAT (red) and vAChT (green) in individual C boutons. (**i**) Despite the conditional knockout of *ChAT* and the complete knockout of *Cart* in *Dbx1::Cre;ChAT*^*fl/fl*^;*CART*^*KO/KO*^ mice, the presynaptic components of C boutons remain intact, as evident by the presence of vAChT (green). (**j**) High magnification image showing the presence of vAChT (green) in individual C boutons that lack both ChAT and CART. (**k**) The ChAT+/vAChT+ C boutons (blue and green respectively) on motor neurons of control *ChAT*^*fl/+*^; *CART*^*KO/+*^ mice are in close apposition to M2 muscarinic receptor clusters (red). (**l**) High magnification image showing the alignment of the post-synaptic M2 muscarinic receptor (red) with the ChAT+/vAChT+ presynaptic part of the synapse (red and green respectively). (**m**) Post synaptic clustering of M2 receptors (red) with presynaptic vAChT (green) is unaltered in *Dbx1::Cre;ChAT*^*fl/fl*^;*CART*^*KO/KO*^ double KO mice. (**n**) High magnification image showing the alignment of M2 postsynaptic muscarinic receptors (red) with the vAChT+ (green) C boutons in *Dbx1*^*cre/+*^*;ChAT*^*fl/fl*^*;CART*^*KO/KO*^ mice. (**o-q**) No change was observed in the number of vAChT+ terminals (p = 0.6730 two-tailed), M2 muscarinic receptor clusters (p = 0.4402 two-tailed) or their alignment (p = 0.2880 two-tailed) despite the simultaneous elimination of ChAT and CART in C boutons. The bar-charts represent the mean number of synapses per motor neuron with superimposed data points representing the data distribution. Data are presented as mean values (±SEM) and were combined as no sex-dependent differences where observed. The median is also presented as a red dot superimposed in each graph. Scale bars = 20μm, (**a, c, e, g, i, k, m**), 1μm (**h, j, l, n**).

We next considered the possibility that CART and acetylcholine may serve parallel roles in supporting C bouton synapses, such that deletion of one could support synapse integrity in the absence of the other. We therefore examined C boutons in double knockout mice (KO), with both global knockout of CART and V0c-selective deletion of ChAT (*Dbx1::Cre;ChAT*^*fl/fl*^*;CART*^*KO/KO*^, double KO; Supplementary Fig. 5a-f). However, examination of the C bouton synapses on the somata of motor neurons of double KO mice using the cholinergic marker vAChT, revealed that the C boutons remain detectable despite the absence of ChAT and CART (Fig. 6g-j). No difference was observed in the number of vAChT+ C boutons in double KO mice compared to WT (Mann-Whitney U = 1867, p = 0.6730 two-tailed; Fig. 6o; Supplementary Table 3) nor in the size of vAChT+ terminals (Mann-Whitney U = 25733, p = 0.0666 two tailed; Supplementary Table 4).

We also considered the possibility that deletion of CART and ChAT could influence the clustering of the key receptor expressed on the postsynaptic membrane at C bouton synapses. We therefore examined the postsynaptic M2 receptors (Fig. 6k-n). We found no difference in either the total number of M2 receptor clusters on motor neuron somata (Mann-Whitney U = 1001, p = 0.4402 two-tailed; Fig. 6p; ST1) nor in the size of M2 receptor clusters (Mann-Whitney U = 25658, p = 0.0537 two tailed; ST2). The number of M2 receptor clusters aligned with vAChT+ boutons also remained unaltered in double knockout mice compared to wildtype controls (Mann-Whitney U = 963, p = 0.2880 two-tailed; Fig. 6q; ST1). Together, these data suggest that neither CART nor acetylcholine are essential for the maintenance of the pre and postsynaptic structure of C boutons, given that we find no change in C bouton number, gross synapse morphology and size, or postsynaptic organization and M2 cluster size in double knockout mice with V0c specific deletion of ChAT and global CART knockout.

### CART and acetylcholine are critical signalling molecules for vigorous motor behaviors

Given that CART is expressed at C boutons and provides amplification of motor neuron output in parallel with acetylcholine mediated signaling, we next set out to determine whether perturbing CART or acetylcholine signaling at C boutons could influence motor behaviors that require greater levels of strength and muscle coordination. We deployed a hanging wire test, which requires mice to recruit multiple muscle groups of the torso and limbs, providing a useful tool to evaluate possible impairments in high muscle force, exertion, and coordination. In this task, mice are positioned on an elevated wire from which they hang with all four limbs. The number of times that animals reached the end of the hanging wire, during a three-minute period, was counted as an indirect measure of muscle strength and endurance ^34–36^. Motor performance on this task was assessed in mice with global CART knockout, C bouton-specific deletion of ChAT, or both manipulations in tandem (double KO).

Adult mice that had ChAT deleted from C boutons (*Dbx1::Cre;ChAT*^*fl/fl*^, n = 22, mean ±SEM = 4.59 ± 0.65) exhibited a significant decrease in the number of times they reached the end of the wire compared to control mice (*ChAT* ^*fl/fl*^ mice, n = 23, mean ±SEM = 7.09 ± 0.90; t (43) = -2.28, p = 0.031). This result indicates that motor performance related to strength is diminished when mice have the ChAT gene exclusively eliminated in V0c interneurons (Fig. 7a). Consistent with animals that had ChAT eliminated from C boutons (*Dbx1::Cre;ChAT*^*fl/fl*^), CART knock out mice (CART^KO/KO^, n =17, mean ±SEM = 5.76 ± 0.70) also reached the end of the wire significantly fewer times compared to wild type control mice, (n= 11, mean ±SEM = 9.27 ± 1.04, t (26) = -2.89, p = 0.008); however, this effect was specific to males only (Fig. 7b). The female CART KO mice (n = 16, mean ±SEM = 9.94 ± 1.56) did not differ significantly in the number of reaches from their wild type counterparts, (n = 11, mean ±SEM = 7.55 ± 1.57, t (25) = 1.04, p = 0.307; Fig. 7c). A similar, significant decrease in the number of times the end of the wire was reached was found in male mice lacking both CART globally and ChAT exclusively from V0c interneurons (*Dbx1::Cre;ChAT*^*fl/fl*^*;CART*^*KO/KO*^, n = 21, mean ±SEM = 3.9 ± 0.50) compared to control mice (n = 28, mean ±SEM = 5.68 ± 0.57, t (47) = -2.25, p = 0.029) (Fig.7d) suggesting a sex-specific motor deficit for this particular genotype. The female counterparts (*Dbx1::Cre;ChAT*^*fl/fl*^*;CART*^*KO/KO*^, n = 15, mean ±SEM = 6.33 ± 0.98) did not differ significantly compared to control mice (n = 25, mean ±SEM = 6.16 ± 0.58, t (38) = 0.163, p = 0.872), Fig. 7e, f).

**Fig. 7:**
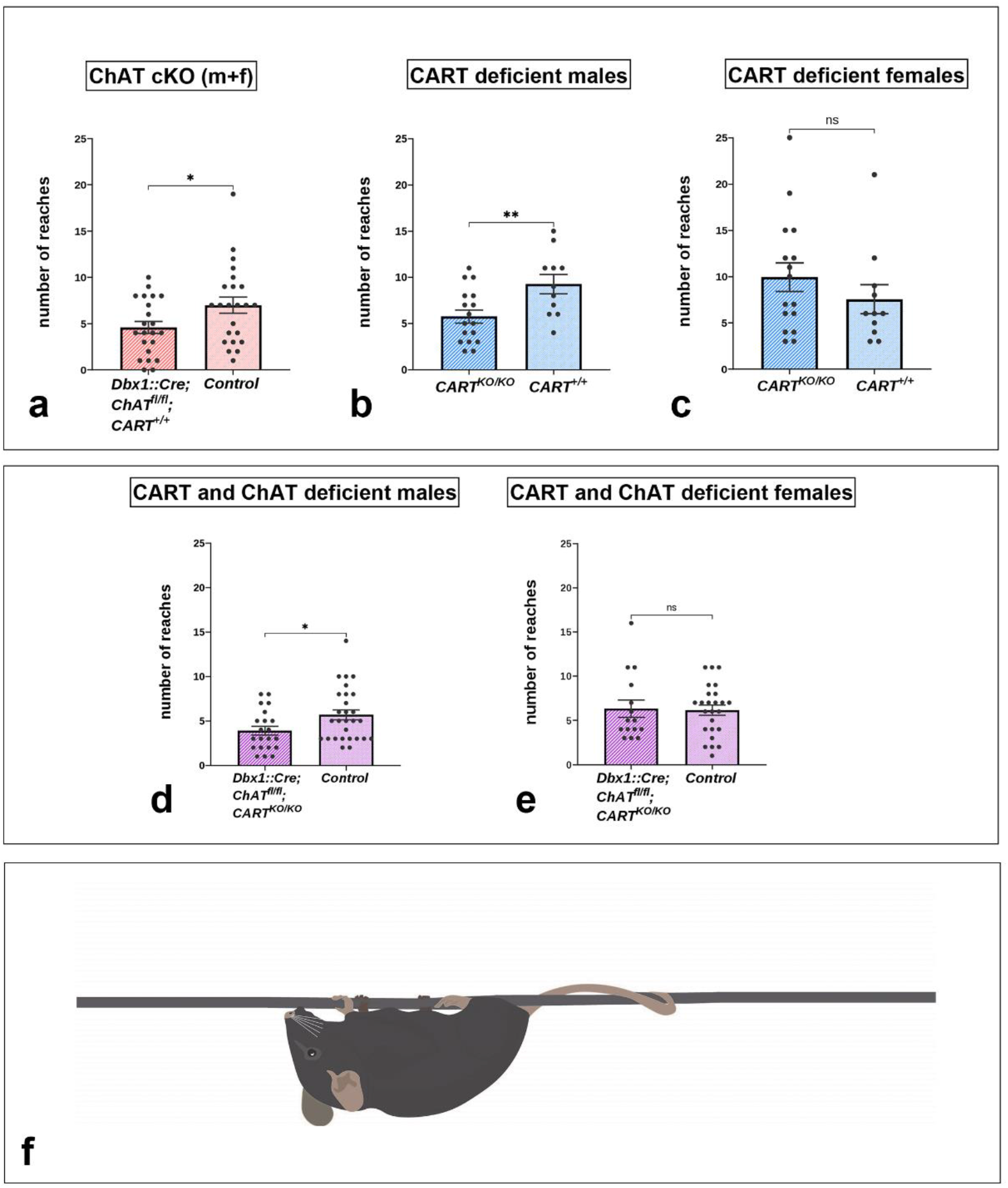
Behavioral effects of the complete knockout of the *Cart* gene and the conditional knockout of the *ChAT* gene in C boutons. Hanging wire test analysis (reaches method) for monitoring muscle strength in mice lacking ChAT in C boutons or CART or of both. Muscle strength is reflected by the number of times that each mouse reaches the end of the wire trying to escape after being suspended at the middle of a 55cm long wire (reaches). (**a**) *Dbx1:Cre;ChAT*^*fl/fl*^ mice (males and females combined) performed significantly fewer reaches than the control *ChAT*^*fl/fl*^ littermates (p = 0.031, two tailed). ***(a)*** *CART KO* male mice performed significantly fewer reaches than the *CART* ^*+/+*^ male mice (p = 0.008, two tailed). (**c**) There were no significant differences between the *CART KO* female mice and the *CART* ^*+/+*^ female mice in the number of reaches performed (p = 0.307, two tailed). (**d**) *Dbx1::Cre;ChAT*^*fl/fl*^*;CART*^*KO/KO*^ male mice performed significantly fewer reaches than the control male mice (p = 0.029, two tailed). (**e**) There were no significant differences between the *Dbx1::Cre;ChAT*^*fl/fl*^*;CART*^*KO/KO*^ female mice and the control female mice in the number of reaches performed (p = 0.872, two tailed). Data are expressed as mean (±SEM). * p < 0.05, ** p < 0.01. (**f**) Illustration of the mice in the starting position of the hanging wire test suspended by all four limbs from the middle of the wire in the behavioral apparatus.

Together these results indicate that CART and acetylcholine exhibit parallel roles in the control of motor behavior, with a particular role for CART in male mice.

## Discussion

Spinal circuits composed of distributed populations of interneurons support a diverse array of functions including the control of movement. Much of what we have learned about roles for spinal interneurons for the control of movement has been derived from the study of locomotor movements that can be broken into properties related to the rhythm and pattern ^2,37,38^. However, complex behaviors are critically dependent on the ability to fine tune the degree of precision and vigour of the underlying movements.

Our previous work identified a subset of spinal cholinergic interneurons, called V0c interneurons that are defined by the expression of the transcription factor Pitx2, and constitute the sole source of C bouton synapses on motor neurons. The C bouton synapse is believed to be an important neuromodulatory synapse that task-dependently increases motor output in mammals ^7,11^. Pre-synaptic components of C boutons (V0c terminals) include cholinergic markers ChAT and vAChT, ^39,40^ along with Synaptobrevin (Vamp2) ^41^, ErbB2/4 receptors ^42^, P2X7R purinergic receptors^43^, and possibly Nicotinic Acetylcholine Receptors (NAchR) ^44^. The post-synaptic membrane aligned with C bouton terminals houses N-type calcium channels ^45^, Ca2+-dependent-K+ channels (SK2/3) ^25^, calcium-activated chloride channels ^26^ and ErbB3/4 receptors ^46^. In the Subsurface Cisterns (SSCs) of C boutons, Sigma-1 receptors ^47^, indole-N-methyl transferase (INMT) ^47^ and Neuregulin 1 (NRG1) ^48^ have been identified (Fig. 2d). The wide range of proteins localized to pre- and post-synaptic regions of C boutons points towards a particularly complicated multicomponent synapse. Here, we have demonstrated further complexity to this synapse, by revealing a novel neuropeptide, encoded by the cocaine and amphetamine regulated transcript (CART) in V0c interneurons at all levels of the spinal cord and in the vast majority of the cholinergic C bouton synapses on motor neurons. These initial anatomical data led us to investigate whether CART may provide a novel complementary signalling mechanism at this important neuromodulatory synapse.

CART was first described in the rat striatum as a hyper-expressed mRNA after cocaine and amphetamine administration ^49^. Since then, CART has been detected throughout the CNS and has been linked to multiple functions, such as olfaction and vision ^19,50^ pain processing ^19,51,52^, addictive behavior ^53,54^, regulation of appetite and body weight ^30,55,56^ as well as stress responses and anxiety ^57–59^. In the spinal cord CART has been previously traced in fibers in laminae I and II of the dorsal horn, in some somata in laminae X and VII, and in sympathetic preganglionic neurons ^19,20,60^.

The fact that we detected CART in synapses on motor neurons raised the question of its involvement in motor control. Knockout of CART exerted functional consequences and led to a poorer performance on the hanging wire test, a motor task that requires increased muscle strength and endurance, pointing to an important role in motor control (Fig. 7). Surprisingly, this deficit was only observed in males. This finding is perhaps not entirely unexpected, given that sex differences regarding CART expression during a stressful motor demanding task have been previously observed ^61^. Also, male specific changes of the C bouton synapse have been observed in disease^62,63^. Although there are currently limited tools (conditional knock out mice, agonists, antagonists) available to study CART’s specific role in motor control, it is possible that this behavioral deficit might be due to the loss of CART in other areas of the CNS. However, our *in vitro* electrophysiological analyses provided evidence that CART influences spinal circuits. Here, we show that CART and acetylcholine, provide differential control of motor neuron subtypes that innervate different muscle fibers; with a bias toward motor neuron subtypes that innervate high force generating, fast twitch muscle fibers that are generally recruited during tasks that require increased levels of muscle force (Fig. 8a-d). Interestingly, CART and acetylcholine accomplish this control through distinct, yet parallel physiological mechanisms for the gradation of muscle force with CART decreasing the recruitment current and acetylcholine agonist muscarine increasing firing rates across much of the frequency current range.

**Fig. 8:**
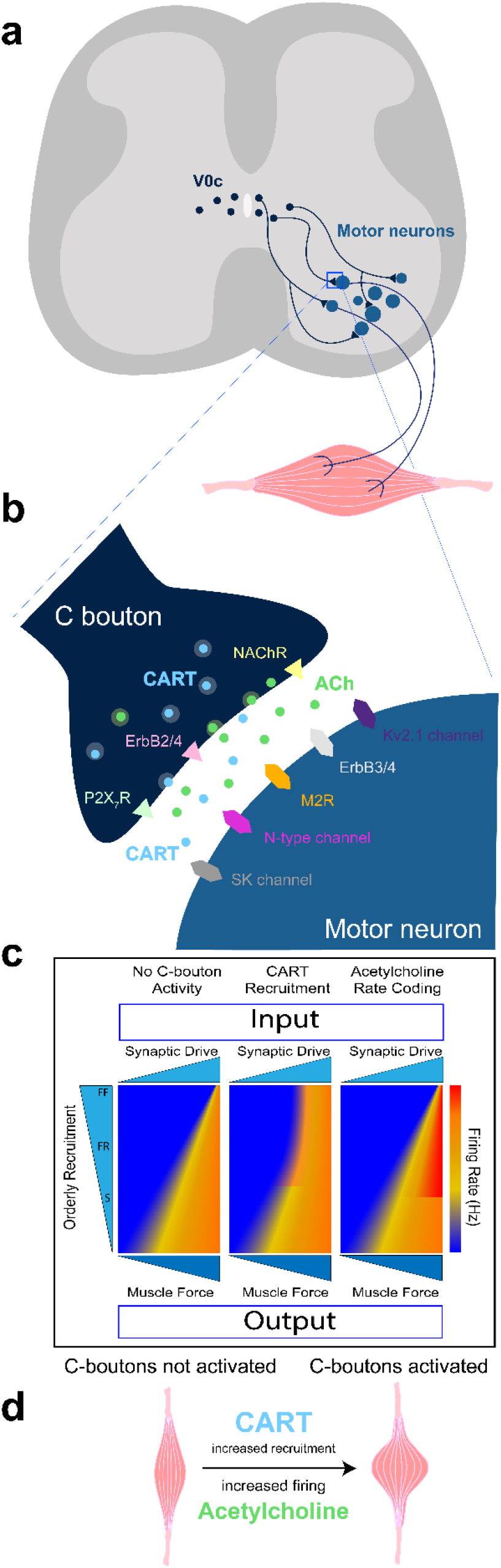
Proposed model of the effect of C bouton function in motor neuron output through acetylcholine and CART signaling. **(a)** Schematic representation of a spinal cord cross section. Dark blue dots represent the V0c somata in the vicinity of the central canal, the axons of which innervate fast and slow motor neuron somata (light blue) in lamina IX (only axons travelling on the same level shown for simplicity). The framed area indicates a single C bouton synapse magnified in B. (**b**) Schematic representation of a single C bouton. The presynaptic terminal (dark blue) contains vesicles loaded with acetylcholine (green) as well as what we propose are Dense Core Vesicles (DCVs) containing the CART peptide (light blue). The presynaptic ATP receptors P2X7R (light green triangles) the ErbB2/4 NRG1 receptors (light pink triangles) and the NAChR - Nicotinic Acetylcholine receptors (light yellow triangles) are also depicted. In the postsynaptic terminal (light blue) SK Ca2+ dependent K+ channels (grey), N-type voltage dependent K+ channels (magenta) and ErbB3/4 NRG1 receptors (light grey) as well as Kv2.1 K+ channels (purple) and M2-muscarinic type 2 acetylcholine receptors- (orange), are presented. (**c**) Schematic representation of how orderly recruitment of slow (S), fast fatigue resistant (FR), and fast fatigable (FF) motor neuron subtypes, and their respective firing rate (heat map) transform synaptic drive (input) to muscle force (output). CART increases output by decreasing the recruitment threshold of fast motor neurons, acetylcholine increases output by increasing fast motor neuron maximal firing rate. (**d**) Suggested effect of C bouton activation in muscle contraction. The combinatorial effect of the CART peptide increasing motor neurons recruitment and acetylcholine increasing the motor neurons’ firing rate leads to a pronounced muscle contraction.

We can now correlate these single cell mechanisms to changes in animal behavior, since we observed that lack of CART in male mice leads to poorer motor performance. It remains to be determined whether peptidergic co-transmission involving CART is a common feature of other cholinergic modulatory systems that regulate the output neurons of a range of neural networks ^13–15^.

The post-synaptic receptors and target ion channels that mediate CART and acetylcholine’s differential control of motor neuron subtypes are not yet known. However, differences in ion channels that control the recruitment and firing rates of fast and slow motor neurons are beginning to be described and could contribute to their differential modulation ^25,28^. Further, recent evidence points to greater C bouton innervation of motor neurons that innervate fast compared to slow twitch muscles; a difference that is most pronounced in male mice^63^, and is in line with behavioral deficits that we observe in CART knockout animals. Regarding CART receptors, recent studies have pinpointed the G-Protein-Coupled-Receptor (GPCR) Gpr160 as a putative receptor for the CART peptide, proposing that CARTp/Gpr160 signaling in the dorsal horn of the spinal cord mediates neuropathic pain through a c-fos and ERK/CREB pathway ^64^. Furthermore, CARTp/Gpr160 coupling in the rat brainstem has been proven necessary for the anorexigenic and antidipsogenic effects of exogenous CART administration ^65^. The identification of the CART peptide receptor in the ventral horn and in particular in the motor neuron membrane remains to be determined; however, its discovery will have the potential to reveal the intracellular signaling mechanisms initiated by CART-motor neuron communication.

C bouton source neurons (Pitx2+ and CART+), the V0c interneurons, were traditionally believed to increase motor output by releasing acetylcholine that acts on M2 muscarinic receptors. Given that chemogenetic activation of V0c interneurons produced only a subset of the effects exerted by global pharmacological activation of muscarinic receptors ^11^, it has been suggested that C boutons produce a distinct set of cholinergic modulatory actions on motor output, with other effects produced by additional, as yet undefined, cholinergic inputs. However, here we show that CART replicates a subset of the effects of chemogenetic activation of V0c interneurons that are not elicited through pharmacological activation of muscarinic receptors, which includes decreased rheobase, increased input resistance, and increased mAHP amplitude. This result suggests that the CART and acetylcholine may provide co-modulation of motor neurons at C boutons synapses. The diversity in modulatory actions between studies could be accounted for by distinct control of fast and slow motor neuron subtypes, which is only beginning to be studied. The diversity in CART function, apart from the recipient motor neuron, is also reflected in the diverse CART expression profiles at C bouton synapses, which were previously believed to be a homogenous synapse. CART is found in most C boutons in different patterns, all of which could be observed throughout the animals’ lifespan (Fig. 3e). In line with our observations, diversity in synapse nanostructure, arising from different inputs, has been reported to also change in ventral versus dorsal areas, during development and through adulthood ^66^. Further, V0c interneuron diversity can also be accounted for by ipsilaterally and bilaterally projecting populations ^67^. C bouton diversity is further supported by the demonstration that V0c interneurons subdivide into connexin36 positive and negative subpopulations ^68^. The gap junction protein is also differentially distributed in C boutons, with fast motor neurons having a greater abundance of connexin 36 positive terminals than slow motor neurons. Thus, even within a small subset of functionally defined spinal interneurons, the V0c subset, we find that there is marked diversity with respect to their anatomy and function depending on source neurons, target motor neuron subtypes, but also sex. This highlights perhaps unexpected but marked diversity in a prominent neuromodulatory synapse, the C bouton synapse, that was once believed to be homogeneous.

The diversity of synaptic components in C bouton synapses, in conjunction with our findings of their functional role, reveal a highly complex and specified component of the motor circuit with a well-defined purpose. Strikingly, we find that the establishment of the V0c-motor neuron synaptic contacts, which are regulated by signals during development, seems to be oblivious of the neurotransmitter content of the C boutons, since the genetic elimination of CART, acetylcholine, or both, did not seem to affect the presence or alter the anatomy of the C bouton synapse nor of its pre and postsynaptic components. This finding lays the ground for the investigation of the wiring of interneuronal motor circuits during development and the elucidation of the importance of functional output for target selection and structural stability of synapses during ontogeny.

In conclusion, we have identified CART neuropeptide together with acetylcholine in the C bouton synapse on motor neurons and proposed a novel role for CART at these synapses, which is important for the neural control of movement. CART facilitates recruitment and firing rates at the lower end of the input range of fast type motor neurons, while acetylcholine augments maximal firing rates throughout most of the input range of fast type motor neurons. Thus, differential neuromodulatory control via acetylcholine and CART may serve as a means to more effectively fine tune muscle activation and force generation. This novel mechanism for adjusting the gain of motor output may be applicable to cholinergic modulatory systems throughout the CNS.

## Methods

### Differential expression screen and in situ hybridization

The differential mRNA expression screen was performed as described in^7^. Briefly upper lumbar (L1-L3) spinal cord segments from P8 mice were isolated and mounted in 2% agarose. The embedded segments were sectioned (Leica VT1000 S Vibrating blade microtome, 200-250μm) and each section was dissected. The intermediate zone was discarded. Dorsal and ventral parts were collected after white matter removal (Fig. 1a). RNA was isolated (n = 3 or 4 mice per sample) using the RNeasy Mini Kit (QIAGEN), and aRNA was synthesized with Ambion’s MessageAmp aRNA Kit (Catalog # 1750) using Biotin 11-CTP and Biotin 16-UTP (Enzo). Affymetrix Gene chip Mouse Genome 430 2.0 Arrays were hybridized. Gene Traffic software was used for data analysis. The *in situ* hybridization experiments for the detection of CART and Pitx2 mRNA expression in spinal cord cross sections (Leica CM3050 S cryostat) were performed as described in ^69^.

### Tissue preparation and immunohistochemistry

Spinal cords were isolated by anesthetizing animals with intraperitoneal injections (15μL/g body weight) of ketamine (10mg/mL) and xylazine (2 mg/mL) prior to performing transcardial perfusion with cold 4% paraformaldehyde (PFA) in PBS (the perfusion was omitted in P0 mice). Spinal cords were isolated and post-fixed in 4% PFA (2h), then washed in PBS (4 times) and cryoprotected in a 30% sucrose solution under slight rocking (4°C) overnight. Spinal cords were further dissected (cervical, thoracic, upper lumbar and lower lumbar segments separation), and embedded in Tissue-Tek OCT medium. The immunohistochemistry experiments were performed in 15-20μm cryostat sections that were collected with positively charged or polylysine covered slides and stored at -80°C after drying in RT. The sections were washed with PBS and were incubated with a 1% BSA-0,1%Triton-X-100 in PBS solution containing the primary antibodies inside a humidified chamber at 4°C overnight. Next the sections were incubated with fluorescently labeled secondary antibodies for 2h, were washed with PBS (4 times for 5 minutes) and covered in Vectashield mounting medium.

Antibodies used: Rabbit anti CART (Phoenix Pharmaceuticals, G003-62), rabbit anti ChAT (Jessell lab, CU1574), rabbit anti vAChT (Jessell lab, CU1475), rabbit anti Pitx2 (Jessell lab, CU1533), rabbit anti M2 (Alomone labs, AB9452), guinea-pig anti vAChT (Fitzerald, 20R-VP002), goat anti vAChT (Millipore, ABN100), goat anti ChAT (Millipore, AB144P).

### Mouse Lines

For all the experiments both male and female animals were utilized. No differences were observed in the anatomy experiments so data were pooled. In the behavioral experiments sex-dependent differences were observed and data were analysed separately. The following mouse lines were used: *Pitx2::Cre* ^21^, *Rosa*.*stop*.*tdTomato* ^22^, *Dbx1::Cre* ^31^, *ChAT* ^*fl/fl* 32,33^, *CART*^*KO/KO* 30^. All genotypes mentioned in this manuscript were a result of elaborate selective breeding of the aforementioned mouse lines. Mice 4-6 months old were used for the behavioral experiments. The use of the specific *CART*^*KO/KO*^ mouse line ^30^ allowed us to perform the motor performance test coherently as the weight gain caused by the complete knockout of the CART gene was detectable only after 40 weeks of age. Anatomy and behavioral experiments were performed in the animal facility of the Center for Clinical, Experimental Surgery and Translational Research of the Biomedical Research Foundation of the Academy of Athens. The facility is registered as “breeding” and “user” establishment by the Veterinary Service of the Prefecture of Athens according to the Presidential Decree 56/2013 in harmonization to the European Directive 63/2010 for the protection of animals used for scientific purposes. User establishment is also ISO 9001:2015 accredited. All animals of the facility were regularly screened using a health monitoring program, in accordance to the Federation of European Laboratory Animal Science Associations’ recommendations.

### Statistical analysis of anatomy data

A total of n=3, *Dbx1::Cre;ChAT*^*fl/fl*^*;CART*^*KO/KO*^ (experimental) and n=4 control mice were used for the synaptic counts. The vAChT+ terminals, postsynaptic M2 receptors and their close apposition were counted form single confocal images that translate to a physical section thickness of 1,208 μm (optical section), using Fiji ^70^. The normality of the distribution of counts within the experimental and control groups was assessed using the Shapiro-Wilk test and the non-parametric Mann-Whitney U test was used to compare group ranks (GraphPad Prism version 9.0.0 for Windows, GraphPad Software, San Diego, California USA, www.graphpad.com). The bar-charts representing the mean number of synapses per motor neuron section with the superimposed data points, representing data distribution, were also designed with GraphPad Prism 9.0.0. The median value is also presented as a red dot superimposed in each graph. Similarly, n=3, *Dbx1::Cre;ChAT*^*fl/fl*^*;CART*^*KO/KO*^ (experimental) and n = 3 control mice were used to evaluate the size of vAChT+ terminals and M2 postsynaptic clusters. In control animals 250 synapses were examined and tested against 228 synapses from experimental animals, using the Mann-Whitney U-test. The same procedure was followed for the 254 control and 225 experimental synapses evaluated for the size of M2 clusters. Data are presented as mean values (±SEM) and were combined as no sex-dependent differences where observed. Statistical significance was set at a < 0.05 for all measures.

### Confocal Imaging and Image analysis using IMARIS (Oxford Instruments)

The 3-Dimensional stack images were captured using a 63x/1.4 NA oil immersion lens and a confocal laser scanning microscope (LeicaTC S SP5 II) in 1024×1024 pixel format. Single images, corresponding to physical thickness of 1,208 μm were used for anatomical analysis and the synapse counting procedures. The z-stack images were used for the reconstruction of the synapse with IMARIS. Using spinal cord sections of *Rosa*.*lsl*.*tdTomato* transgenic mice incubated with antibodies against dsRed and CART the colocalization of the CART peptide in C bouton synapses was visualized. First, confocal images were deconvolved using LAS AF (Leica Microsystems – Blind Deconvolution option) in order to reduce out of focus fluorescence and improve contrast allowing for the accurate reconstruction of < 1 μm structures. Using the surface rendering module of IMARIS, two different volumes were created (3D surfaces) based on the fluorescence intensity and size (red: C bouton synapse; green: CART peptide). The overlap of the surfaces and the enclosure of the CART peptide surface (green) in the C bouton surface (red) was visualized by rendering the synapse exterior (red surface) transparent.

### Tissue preparation for electrophysiology

Experiments were performed on male and female wild type C57 Black J/6 mice from postnatal days 7-12 (n= 30). All procedures were approved by the Home Office and the University of St. Andrews Animal Care Committee. All animals were sacrificed by performing a cervical dislocation followed by rapid decapitation. Animals were eviscerated and pinned ventral side up in a dissecting chamber lined with silicone elastomer (Sylguard) filled carbogenated (95% oxygen, 5% carbon dioxide), pre-chilled (1-4 degrees Celsius) dissecting artificial cerebrospinal fluid (aCSF; in mM: 25 NaCl, 188 sucrose, 1.9 KCl, 1.2 NaH2PO4, 10 MgSO4, 1 CaCl, 26 NaHCO3, 25 D-glucose, 1.5 kynurenic acid). Spinal cords were exposed by performing a ventral laminectomy, ventral roots cut and gently lifted from the spinal column. The lumbar section of the spinal cord was then secured with the ventral surface facing out to an agar block with surgical glue (Vetbond). The agar block was secured to the base of the slicing chamber with super glue. Slices from rostral (L1-L3) or caudal (L4-L6) segments were obtained on different days by placing the rostral or caudal segments facing up. The slicing chamber was filled with pre-chilled (1 - 4 degrees Celsius) carbogenated dissecting aCSF and 300 μm transverse slices made with a cutting speed of 100 μm/s on a Leica Vibratome (Leica, CT1200). Slices were transferred to a recovery chamber filled with carbogenated pre-warmed (35 degrees Celsius) recovery aCSF (in mM: 119 NaCl, 1.9 KCl, 1.2 NaH2PO4, 10 MgSO4, 1 CaCl, 26 NaHCO3, 20 glucose, 1.5 kynurenic acid, 3% dextran) for thirty minutes after completion of the last slice. Following recovery, slices were transferred to a chamber filled with carbogenated warm (35 degrees Celsius) recording aCSF (in mM: 127 NaCl, 3 KCl, 1.25 NaH2PO4, 1 MgCl, 2 CaCl2, 26 NaHCO3, 10 glucose) which was then allowed to equilibrate at room temperature (23-25 degrees Celsius) for at least one hour before experiments were initiated.

### Whole cell patch clamp electrophysiology

Whole cell patch clamp recordings were obtained from motor neurons of rostral and caudal lumbar segments. Slices were stabilized in a recording chamber with fine fibers secured to a platinum harp and visualized on an Olympus BX51WI microscope under a 40x objective with infrared illumination and differential interference (DIC) contrast. Whole cell recordings were obtained under DIC illumination with pipettes (L: 100 mm, OD: 1.5 mm, ID: 0.84 mm; World Precision Instruments) pulled on a Flaming Brown micropipette puller (Sutter instruments P97) to a resistance of 3-4 MOhms. Pipettes were back-filled with intracellular solution (in mM: 140 KMeSO4, 10 NaCl, 1 CaCl2, 10 HEPES, 1 EGTA, 3 Mg-ATP and 0.4 GTP-Na2; pH 7.2-7.3, adjusted with KOH), attached to an electrode holder and preamplifier (Molecular Devices CV-7B), which were positioned with a Patch Star micromanipulator (Scientifica, UK). Signals were amplified and filtered (6 kHz low pass Bessel filter) with a Multiclamp 700 B amplifier, acquired at 20 kHz using a Digidata 1440A digitizer with pClamp Version 10.7 software (Molecular Devices) and stored on a computer for offline analysis. Current clamp recordings were whole-cell capacitance compensated using Multiclamp Commander software.

Motor neurons were identified based on location in the ventrolateral region with somata greater than 20 μm. Intrinsic properties were only measured if cells had an initial resting membrane potential below -50 mV or if access resistance was less than 20 MΩ. All recordings were performed on cells in spinal cord slices that were naive to drug treatment with no more than one cell studied per slice. Cells were excluded from analysis if access resistance changed by more than 20% of the initial value or if action potential amplitude was less than 60 mV measured from threshold. Motor neuron intrinsic properties were studied by applying a bias current to maintain the membrane potential at -60mV. All values reported are not liquid junction potential corrected to facilitate comparisons with previously published data.

### Identification of fast and slow motor neuron types

Fast and slow motor neurons were identified as by ^27,71^ based on the onset latency at repetitive firing threshold using a 5 second square depolarizing current at rheobase. Using this approach, we were able to identify 2 repetitive firing profiles - a delayed repetitive firing with accelerating spike frequency characteristic of fast-type motor neurons and an immediate firing profile characteristic of slow type motor neurons ^27,28,71^.

### Pharmacology

The cocaine and amphetamine regulated transcript peptide (CART acetate salt; 48 Amino Acid sequence: IPIYEKKYGQVPMCDAGEQCAVRKGARIGKLCDCPRGTSCNSFLLKCL; with triple disulfide bonds at Cys14 - Cys32, Cys20 - Cys40, Cys34 - Cys47; Lot No. P170320-MJ470307; MW = 5259.1 g/mol; Cat no.: 470307; Biomatik) and muscarine (Sigma-Aldrich) were dissolved in H_2_O and stored at -20 degrees Celsius until required for experiments. A working concentration of 1 μM CART and 10 μM muscarine were used for in vitro experiments.

### Assessment of recruitment properties

Recruitment properties of motor neurons subtypes were assessed during slow (100 pA/s) depolarizing current ramps which allowed us to measure the recruitment current and voltage threshold of the first action potential. The voltage threshold was defined when the change in voltage reached 10 mV/s. Firing rates were compared at three points on the frequency-current curve: at recruitment (minimum firing rate), 2 times recruitment current, and maximum firing rates.

### Assessment of single action potential (spike) properties

Single spike properties (spike threshold, amplitude, rise time, half width) and AHP properties (amplitude and half width) were assessed with a 10 ms square pulse applied at an intensity 25% above rheobase. Spike threshold was determined as the potential at which the derivative (dV/dt) increased above 10 mV/s. Spike amplitude, rise time, and half width were measured from this point. AHP properties (amplitude and half width) were measured from baseline (−60 mV). Single spike and AHP properties were measured using event detection features in Clampfit.

### Electrophysiology statistical analysis

Mixed-effects ANOVA were conducted to test the effect of pharmacological agents on intrinsic properties with a Geisser-Greenhouse’s correction factor. Bonferroni post hoc analysis was performed when significant main effects were detected with a significance level of p < 0.05. Data are presented in figures as individual data points for all cells. Statistical analyses were performed using Graph Pad Version 9.0.0 (GraphPad Prism version 9.0.0 for Windows, GraphPad Software, San Diego, California USA, www.graphpad.com).

### Hanging wire behavioral test and data analysis

The hanging wire test (reaches method) was used to assess muscle strength and coordination as was adapted from the Treat-NMD Neuromuscular Network standard operating protocol (Van Putten, 2011 DMD_M.2.1.004). Each mouse was handled by the experimenter for five consecutive days before the behavioral task. Mice were moved to the testing room 20-30 min prior to testing. All procedures and testing took place between 13:00-17:00 p.m. in an isolated testing room. A non-bending metallic wire (55cm long, 2.5mm thick) was attached to two transparent Plexiglas surfaces (40cm high) at a height approximately 35cm above soft bedding needed to break possible falls of the mouse. Each mouse handled by the tail by the experimenter was placed to the middle of the wire until it grasped it with all four limbs. For a period of 180sec the experimenter measured how many times the mouse would reach the one or the other end of the wire. Each time the mouse reached the end of the wire (1cm or less from the Plexiglas surface) the timer was stopped and the experimenter positioned the mouse at the middle of the wire hanging with all four limbs and facing the opposite Plexiglas surface. The timer would start again at this point. The number of reaches for each mouse during a three minutes period was used for statistical analysis. Statistical analyses were performed using SPSS Version 22 (SPSS Inc, Chicago, IL, USA). The groups were compared using the independent samples t-test. Data from male and female mice were combined when there were no significant differences between them, or reported separately when they differed significantly. Four to six months old mice were used. Data are presented as mean values (±SEM). Statistical significance was set at a < 0.05 for all measures.

## Acknowledgments

We would like to acknowledge the following people for their contributions to this manuscript: Dr. Niels Wierup for kindly providing us with the *CART*^*KO/KO*^ mouse line, Eirini Eleftheriadi for helping with the graphic design in illustrations of the figures, Despoina Bosveli for further analyzing part of the anatomy data, Dr. Nikolaos Kostomitsopoulos for advice and support at the BRFAA mouse facility and Dr. David L. McLean, Dr. Katharina Quinlan and Dr. Stamatis N. Pagakis for critically reading the manuscript. We would also like to thank BRFAA Imaging Unit personnel for support on imaging.

This work was supported by Greek national funds (ARISTEIA II 4257, LZ), Fondation SANTE (LZ), a Marie Curie Re-Integration Grant (268323, LZ) and by a St Andrews Restarting Research Fund (GBM and SAS). SAS was funded by a Royal Society Newton International Fellowship (NIF/R1/180091) and a Natural Sciences and Engineering Research Council of Canada (NSERC) Postdoctoral Fellowship (NSERC-PDF-517295-2018) and MM by the National Scholarship Foundation (IKY).

## Author contributions

L.Z., G.B.M., P.E.E., S.A.S. and K.P. were involved in conceptualization. L.Z., P.E.E., P.E.A., E.T. and M.M. performed and analyzed anatomy and imaging experiments. S.A.S. performed and analyzed electrophysiology experiments. K.P. performed and analyzed behavioral experiments. L.Z., G.B.M., P.E.E, S.A.S and K.P wrote the manuscript, which was edited by all the authors. L.Z and G.B.M. supervised the project.

## Data availability

The data that support the findings of this study are available from the corresponding author upon reasonable request.

## References

1. Gosgnach, S. et al. Delineating the diversity of spinal interneurons in locomotor circuits. Journal of Neuroscience 37, 10835–10841 (2017).

2. Goulding, M. Circuits controlling vertebrate locomotion: Moving in a new direction. Nat Rev Neurosci 10, 507–518 (2009).

3. Henneman, E. Relation between size of neurons and their susceptibility to discharge. Science (1979) 126, 1345–1347 (1957).

4. Burke, R. E., Levine, D. N., Tsairis, P. & Zajac, F. E. Physiological types and histochemical profiles in motor units of the cat gastrocnemius. J Physiol 234, 723–748 (1973).

5. Milner-Brown, H. S., Stein, R. B. & Yemm, R. Changes in firing rate of human motor units during linearly changing voluntary contractions. J Physiol 230, 371 (1973).

6. Conradi S, Skoglund S. Observations on the ultrastruture and distribution of neuronal and glial elements on the motoneuron surface in the lumbosacral spinal cord of the cat during postnatal development. Acta Physiol Scand Suppl 333, 5–52 (1969).

7. Zagoraiou, L. et al. A cluster of cholinergic premotor interneurons modulates mouse locomotor activity. Neuron 64, 645–662 (2009).

8. Lanuza, G. M., Gosgnach, S., Pierani, A., Jessell, T. M. & Goulding, M. Genetic identification of spinal interneurons that coordinate left-right locomotor activity necessary for walking movements. Neuron 42, 375–386 (2004).

9. Pierani, A. et al. Control of interneuron fate in the developing spinal cord by the progenitor homeodomain protein Dbx1. Neuron 29, 367–384 (2001).

10. Moran-Rivard, L. et al. Evx1 is a postmitotic determinant of V0 interneuron identity in the spinal cord. Neuron 29, 385–399 (2001).

11. Nascimento, F. et al. Synaptic mechanisms underlying modulation of locomotor-related motoneuron output by premotor cholinergic interneurons. Elife 9, (2020).

12. Clamann, H. P. Motor Unit Recruitment and the Gradation of Muscle Force. Phys Ther 73, 830–843 (1993).

13. Xiao, C. et al. Cholinergic Mesopontine Signals Govern Locomotion and Reward through Dissociable Midbrain Pathways. Neuron 90, 333–347 (2016).

14. Pickford, J., Apps, R. & Bashir, Z. I. Muscarinic Receptor Modulation of the Cerebellar Interpositus Nucleus In Vitro. Neurochem Res 44, 627–635 (2019).

15. Li, Y. & Hollis, E. Basal Forebrain Cholinergic Neurons Selectively Drive Coordinated Motor Learning in Mice. J Neurosci 41, 10148–10160 (2021).

16. Kiehn, O. & Butt, S. J. B. Physiological, anatomical and genetic identification of CPG neurons in the developing mammalian spinal cord. Prog Neurobiol 70, 347–361 (2003).

17. Dougherty, K. J. et al. Locomotor Rhythm Generation Linked to the Output of Spinal Shox2 Excitatory Interneurons. Neuron 80, 920–933 (2013).

18. Fenwick, N. M., Martin, C. L. & Llewellyn-Smith, I. J. Immunoreactivity for cocaine- and amphetamine-regulated transcript in rat sympathetic preganglionic neurons projecting to sympathetic ganglia and the adrenal medulla. J Comp Neurol 495, 422–433 (2006).

19. Koylu, E. O., Couceyro, R., Lambert, P. D. & Kuhar, M. J. Cocaine-and Amphetamine-Regulated Transcript Peptide Immunohistochemical Localization in the Rat Brain. J. Comp. Neurol 391, 115–132 (1998).

20. Dun, S. L., Chianca, D. A., Dun, N. J., Yang, J. & Chang, J. K. Differential expression of cocaine- and amphetamine-regulated transcript-immunoreactivity in the rat spinal preganglionic nuclei. Neurosci Lett 294, 143–146 (2000).

21. Liu, W., Selever, J., Lu, M. F. & Martin, J. F. Genetic dissection of Pitx2 in craniofacial development uncovers new functions in branchial arch morphogenes, late aspects of tooth morphogenesis and cell migration. Development 130, 6375–6385 (2003).

22. Madisen, L. et al. A robust and high-throughput Cre reporting and characterization system for the whole mouse brain. Nature Neuroscience 2009 13:1 13, 133–140 (2009).

23. Miles, G. B., Hartley, R., Todd, A. J. & Brownstone, R. M. Spinal cholinergic interneurons regulate the excitability of motoneurons during locomotion. Proc Natl Acad Sci U S A 104, 2448–2453 (2007).

24. Hellström, J., Oliveira, A. L. R., Meister, B. & Cullheim, S. Large cholinergic nerve terminals on subsets of motoneurons and their relation to muscarinic receptor type 2. Journal of Comparative Neurology 460, 476–486 (2003).

25. Deardorff, A. S. et al. Expression of postsynaptic Ca2+-activated K+ (SK) channels at C-bouton synapses in mammalian lumbar α-motoneurons. Journal of Physiology 591, 875–897 (2013).

26. Soulard, C. et al. Spinal Motoneuron TMEM16F Acts at C-boutons to Modulate Motor Resistance and Contributes to ALS Pathogenesis. Cell Rep 30, 2581-2593.e7 (2020).

27. Leroy, F., Lamotte d’Incamps, B., Imhoff-Manuel, R. D. & Zytnicki, D. Early intrinsic hyperexcitability does not contribute to motoneuron degeneration in amyotrophic lateral sclerosis. Elife 3, (2014).

28. Sharples, S. A. & Miles, G. B. Maturation of persistent and hyperpolarization-activated inward currents shapes the differential activation of motoneuron subtypes during postnatal development. Elife 10, (2021).

29. Nascimento, F., Spindler, L. R. B. & Miles, G. B. Balanced cholinergic modulation of spinal locomotor circuits via M2 and M3 muscarinic receptors. Sci Rep 9, (2019).

30. Wierup, N. et al. CART knock out mice have impaired insulin secretion and glucose intolerance, altered beta cell morphology and increased body weight. Regul Pept 129, 203–211 (2005).

31. Bielle, F. et al. Multiple origins of Cajal-Retzius cells at the borders of the developing pallium. Nat Neurosci 8, 1002–1012 (2005).

32. Misgeld, T. et al. Roles of neurotransmitter in synapse formation: development of neuromuscular junctions lacking choline acetyltransferase. Neuron 36, 635–648 (2002).

33. Buffelli, M. et al. Genetic evidence that relative synaptic efficacy biases the outcome of synaptic competition. Nature 424, 430–434 (2003).

34. Kundu, P. et al. Integrated analysis of behavioral, epigenetic, and gut microbiome analyses in App NL-G-F, App NL-F, and wild type mice. Sci Rep 11, (2021).

35. Martinez-Huenchullan, S. F. et al. Utility and reliability of non-invasive muscle function tests in high-fat-fed mice. Exp Physiol 102, 773–778 (2017).

36. Luna-Sánchez, M. et al. The clinical heterogeneity of coenzyme Q10 deficiency results from genotypic differences in the Coq9 gene. EMBO Mol Med 7, 670–687 (2015).

37. Grillner, S. & el Manira, A. Current Principles of Motor Control, with Special Reference to Vertebrate Locomotion. Physiol Rev 100, 271–320 (2020).

38. Kiehn, O. Decoding the organization of spinal circuits that control locomotion. Nat Rev Neurosci 17, 224–238 (2016).

39. Connaughton, M., Priestley, J. v., Sofroniew, M. v., Eckenstein, F. & Cuello, A. C. Inputs to motoneurones in the hypoglossal nucleus of the rat: Light and electron microscopic immunocytochemistry for choline acetyltransferase, substance P and enkephalins using monoclonal antibodies. Neuroscience 17, 205–224 (1986).

40. Nachmansohn, D. & Machado, A. L. THE FORMATION OF ACETYLCHOLINE. A NEW ENZYME: ‘CHOLINE ACETYLASE’. J Neurophysiol 6, 397–403 (1943).

41. Hellström, J., Arvidsson, U., Elde, R., Cullheim, S. & Meister, B. Differential expression of nerve terminal protein isoforms in VAChT-containing varicosities of the spinal cord ventral horn. J Comp Neurol 411, 578–590 (1999).

42. Casanovas, A. et al. Neuregulin 1-ErbB module in C-bouton synapses on somatic motor neurons: Molecular compartmentation and response to peripheral nerve injury. Sci Rep 7, 1–17 (2017).

43. Deng, Z. & Fyffe, R. E. W. Expression of P2X7 receptor immunoreactivity in distinct subsets of synaptic terminals in the ventral horn of rat lumbar spinal cord. Brain Res 1020, 53–61 (2004).

44. Khan, I. et al. Nicotinic acetylcholine receptor distribution in relation to spinal neurotransmission pathways. Journal of Comparative Neurology 467, 44–59 (2003).

45. Wilson, J. M., Rempel, J. & Brownstone, R. M. Postnatal development of cholinergic synapses on mouse spinal motoneurons. J Comp Neurol 474, 13–23 (2004).

46. Lasiene, J. et al. Neuregulin 1 confers neuroprotection in SOD1-linked amyotrophic lateral sclerosis mice via restoration of C-boutons of spinal motor neurons. Acta Neuropathol Commun 4, 15 (2016).

47. Mavlyutov, T. A. et al. Development of the sigma-1 receptor in C-terminals of motoneurons and colocalization with the N,N’-dimethyltryptamine forming enzyme, indole-N-methyl transferase. Neuroscience 206, 60–68 (2012).

48. Gallart-Palau, X. et al. Neuregulin-1 is concentrated in the postsynaptic subsurface cistern of C-bouton inputs to α-motoneurons and altered during motoneuron diseases. The FASEB Journal 28, 3618–3632 (2014).

49. Douglass, J., McKinzie, A. A. & Couceyro, P. PCR differential display identifies a rat brain mRNA that is transcriptionally regulated by cocaine and amphetamine. Journal of Neuroscience 15, 2471–2481 (1995).

50. Couceyro, P. R., Koylu, E. O. & Kuhar, M. J. Further studies on the anatomical distribution of CART by in situ hybridization. J Chem Neuroanat 12, 229–241 (1997).

51. Imad Damaj, M., Zheng, J., Martin, B. R. & Kuhar, M. J. Intrathecal CART (55-102) attenuates hyperlagesia and allodynia in a mouse model of neuropathic but not inflammatory pain. Peptides (N.Y.) 27, 2019–2023 (2006).

52. Ohsawa, M., Dun, S. L., Tseng, L. F., Chang, J. K. & Dun, N. J. Decrease of hindpaw withdrawal latency by cocaine-and amphetamine-regulated transcript peptide to the mouse spinal cord. Eur J Pharmacol 399, 165–169 (2000).

53. Jaworski, J. N., Vicentic, A., Hunter, R. G., Kimmel, H. L. & Kuhar, M. J. CART peptides are modulators of mesolimbic dopamine and psychostimulants. Life Sci 73, 741–747 (2003).

54. Lohoff, F. W. et al. Genetic variants in the cocaine- and amphetamine-regulated transcript gene (CARTPT) and cocaine dependence. Neurosci Lett 440, 280–283 (2008).

55. de Lartigue, G. et al. Cocaine- and Amphetamine-Regulated Transcript Mediates the Actions of Cholecystokinin on Rat Vagal Afferent Neurons. Gastroenterology 138, 1479–1490 (2010).

56. Ludvigsen, S., Thim, L., Blom, A. M. & Wulff, B. S. Solution structure of the satiety factor, CART, reveals new functionality of a well-known fold. Biochemistry 40, 9082–9088 (2001).

57. Koylu, E. O., Balkan, B., Kuhar, M. J. & Pogun, S. Cocaine and amphetamine regulated transcript (CART) and the stress response. Peptides vol. 27 1956–1969 Preprint at https://doi.org/10.1016/j.peptides.2006.03.032 (2006).

58. Smith, S. M. et al. Cocaine- and amphetamine-regulated transcript activates the hypothalamic-pituitary-adrenal axis through a corticotropin-releasing factor receptor-dependent mechanism. Endocrinology 145, 5202–5209 (2004).

59. Dandekar, M. P. et al. Importance of cocaine- and amphetamine-regulated transcript peptide in the central nucleus of amygdala in anxiogenic responses induced by ethanol withdrawal. Neuropsychopharmacology 33, 1127–1136 (2008).

60. Dun, S. L., Ng, Y. K., Brailoiu, G. C., Ling, E. A. & Dun, N. J. Cocaine- and amphetamine-regulated transcript peptide-immunoreactivity in adrenergic C1 neurons projecting to the intermediolateral cell column of the rat. J Chem Neuroanat 23, 123–132 (2002).

61. Balkan, B. et al. CART expression in limbic regions of rat brain following forced swim stress: Sex differences. Neuropeptides 40, 185–193 (2006).

62. Herron, L. R. & Miles, G. B. Gender-specific perturbations in modulatory inputs to motoneurons in a mouse model of amyotrophic lateral sclerosis. Neuroscience 226, 313–323 (2012).

63. Bak, A. N. et al. Cytoplasmic TDP-43 accumulation drives changes in C-bouton number and size in a mouse model of sporadic Amyotrophic Lateral Sclerosis. bioRxiv 2022.05.20.492885 (2022) doi:10.1101/2022.05.20.492885.

64. Yosten, G. L. C. et al. GPR160 de-orphanization reveals critical roles in neuropathic pain in rodents. J Clin Invest 130, 2587 (2020).

65. Haddock, C. J. et al. Signaling in rat brainstem via Gpr160 is required for the anorexigenic and antidipsogenic actions of cocaine- and amphetamine-regulated transcript peptide. https://doi.org/10.1152/ajpregu.00096.2020 320, R236–R249 (2021).

66. Broadhead, M. J. et al. Nanostructural Diversity of Synapses in the Mammalian Spinal Cord. Scientific Reports \x2020 10:1 10, 1–18 (2020).

67. Stepien, A. E., Tripodi, M. & Arber, S. Monosynaptic Rabies Virus Reveals Premotor Network Organization and Synaptic Specificity of Cholinergic Partition Cells. Neuron 68, 456–472 (2010).

68. Recabal-Beyer, A. J., Senecal, J. M. M., Senecal, J. E. M., Lynn, B. D. & Nagy, J. I. On the Organization of Connexin36 Expression in Electrically Coupled Cholinergic V0c Neurons (Partition Cells) in the Spinal Cord and Their C-terminal Innervation of Motoneurons. Neuroscience 485, 91–115 (2022).

69. Schaeren-Wiemers, N. & Gerfin-Moser, A. A single protocol to detect transcripts of various types and expression levels in neural tissue and cultured cells: in situ hybridization using digoxigenin-labelled cRNA probes. Histochemistry 100, 431–440 (1993).

70. Schindelin, J. et al. Fiji: an open-source platform for biological-image analysis. Nature Methods 2012 9:7 9, 676–682 (2012).

71. Leroy, F. & Zytnicki, D. Is hyperexcitability really guilty in amyotrophic lateral sclerosis? Neural Regen Res 10, 1413 (2015).

