## Supplementary Information for "“Peptidergic modulation of motor neuron output via CART signaling at C bouton synapses”"

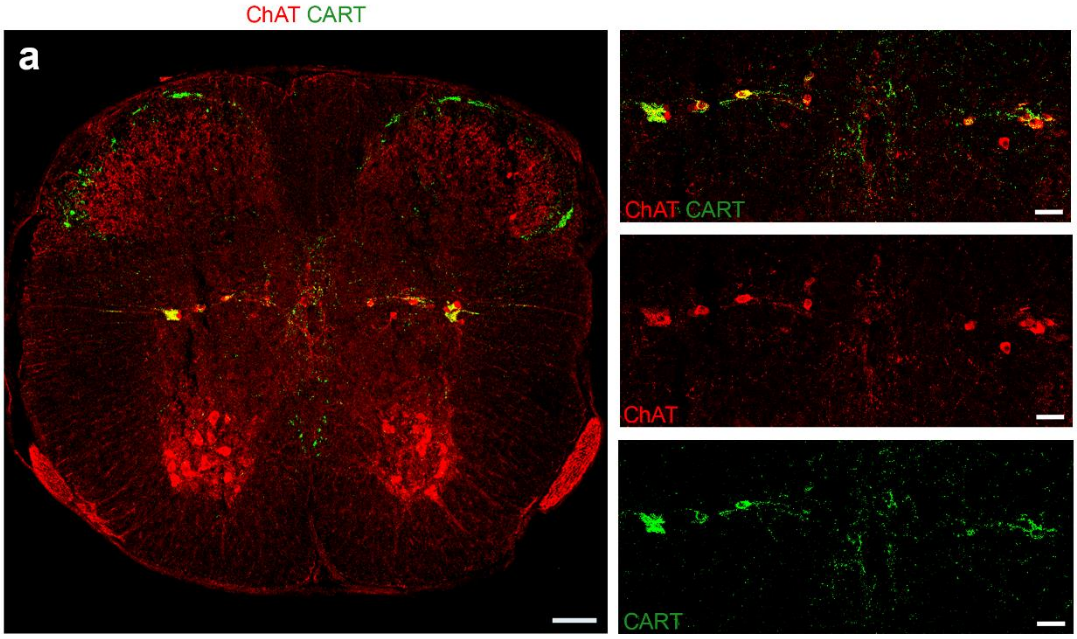


**Supplementary Fig. 1: Sympathetic preganglionic neurons express CART**

(**a**) Immunofluorescence in thoracic spinal cord cross section of P25 mice with antibodies against CART (green) and ChAT (red). Sympathetic preganglionic cholinergic neurons express the CART peptide, reaffirming previous reports. Higher magnification panels focusing on the intermediate zone show merged and separate channels. Scale bars: (**a**) 100mµ (panels) 20µm.

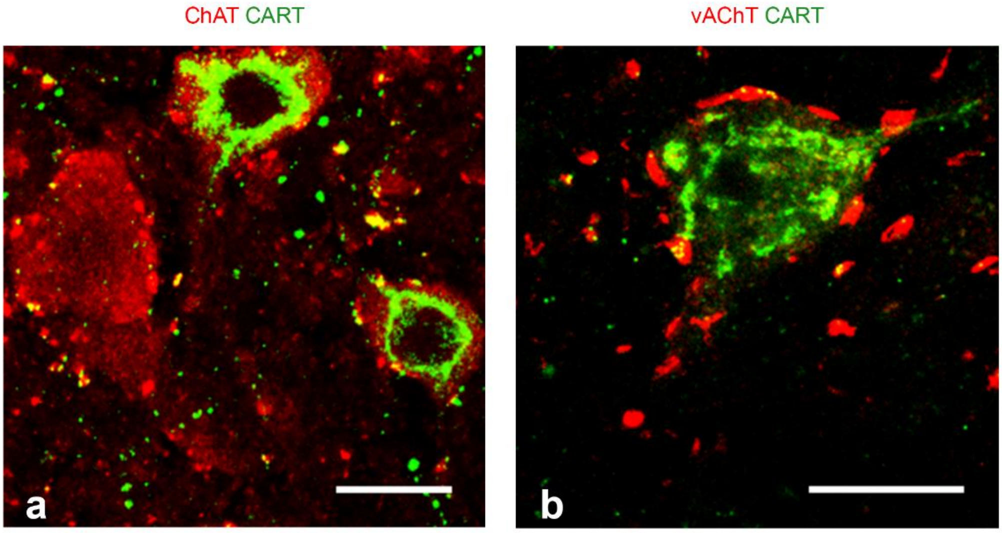


**Supplementary Fig. 2: A subset of motor neurons also expresses the CART neuropeptide**

(**a,b**) Immunofluorescence for ChAT (red) and CART (green) reveals CART immunoreactivity in the somata of few motor neurons as well as CART+ C boutons on CART+ motor neurons. Scale bar: 20 µm

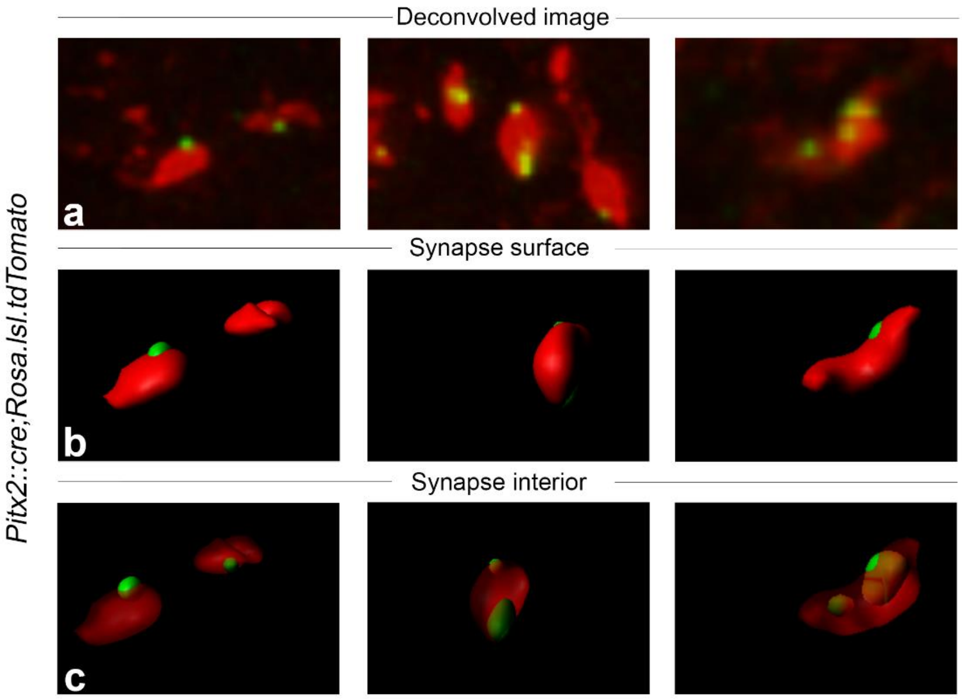


**Supplementary Fig. 3: 3D reconstruction of the C bouton synapse containing the CART neuropeptide**

IMARIS software-based 3D reconstruction of the synapse volume from confocal images of *Pitx2::Cre;Rosa.lsl.tdTomato* spinal cord sections with antibodies against dsRed (red) and CART (green). (**a**) Deconvolved z-stack images of the C bouton synapses used for the reconstruction. (**b**) The synapse volume is reconstructed in 3D. (**c**) The synapse is rendered transparent revealing the distribution of CART in the interior of the synapse.

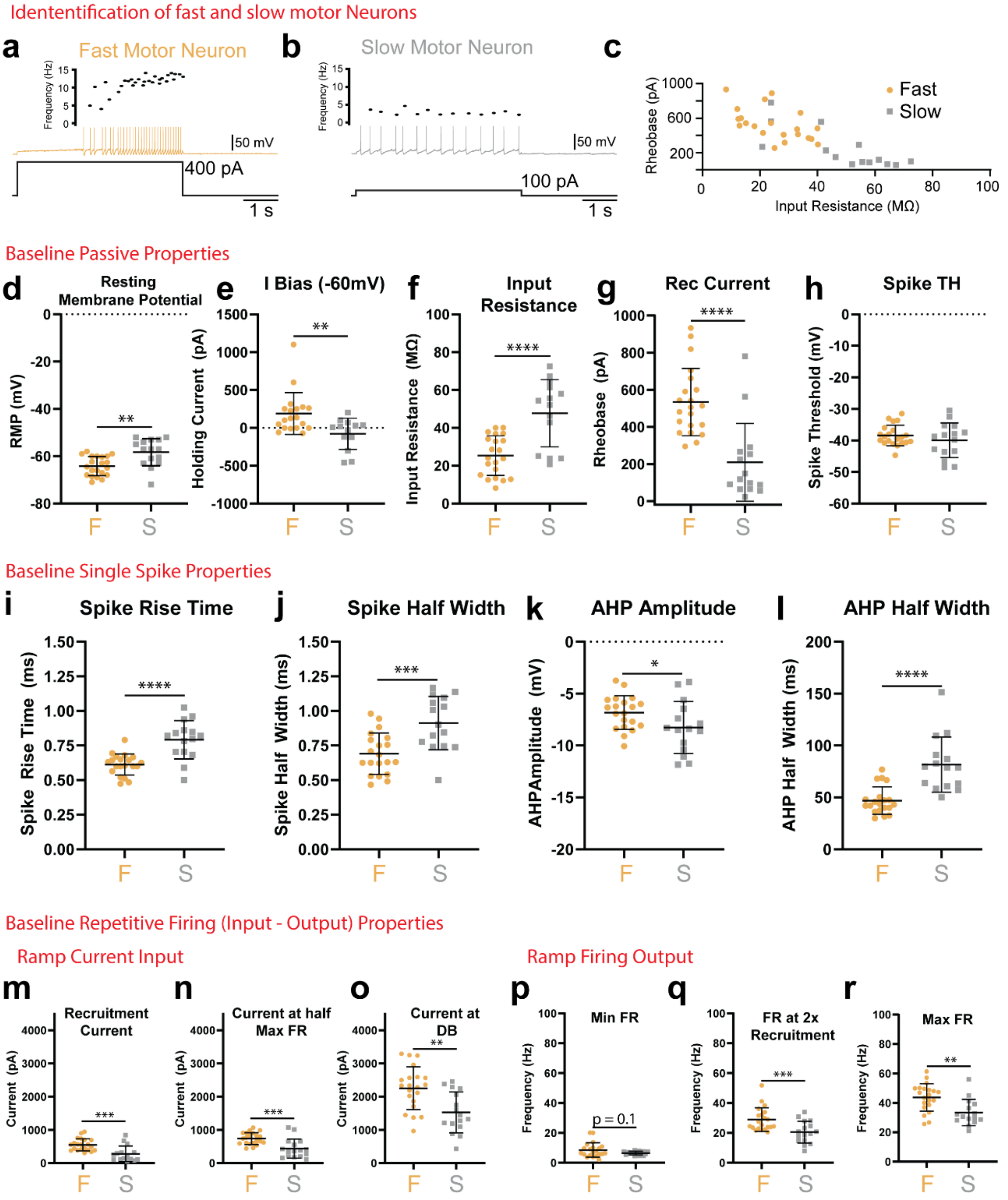


**Supplementary Fig. 4: Properties of fast and slow type motor neurons at baseline**

(**a, b**) Representative traces of fast (**a**: Orange) and slow (**b**: grey) motor neurons identified with whole cell patch clamp electrophysiology which display delayed or immediate repetitive firing profiles, respectively. (**c**) Scatter plot between input resistance and recruitment current (rheobase) demonstrate that fast motor neurons have a high rheobase, low input resistance, whereas slow motor neurons have a low rheobase and high input resistance. Compared to slow motor neurons, fast motor neurons have a significantly more hyperpolarized resting membrane potential (**d**), require more bias current to bring the RMP to - 60 mV (**e**), have a lower input resistance (**f**), higher recruitment current (**g**), with no difference in spike threshold (**h**). Single action potential (spikes) features of fast motor neurons are significantly faster, with a shorter spike rise time (**i**) and half width (**j**). The medium afterhyperpolarization (mAHP) is significantly smaller in amplitude (**k**) and shorter half width (**l**) in fast compared to slow motor neurons. The frequency-current (input-output) relationship was assessed on a slow depolarizing current ramp (100 pA/s). Input values (Currents) at repetitive firing onset (**m**; recruitment), half maximum firing (**n**) and depolarizing block (DB; **o**) were significantly higher in fast compared to slow motor neurons. The output (firing rate) was not significantly different at recruitment (**p**; p=0.1) but was significantly higher in fast compared to slow motor neurons at 2x recruitment current (**q**) and maximal firing rates (**r**). Horizontal black lines depict the mean and error bars represent standard deviation. Asterisks denote significance (*p<0.05, **p<0.01, ***p<0.001. ****p<0.0001) from unpaired t-test.

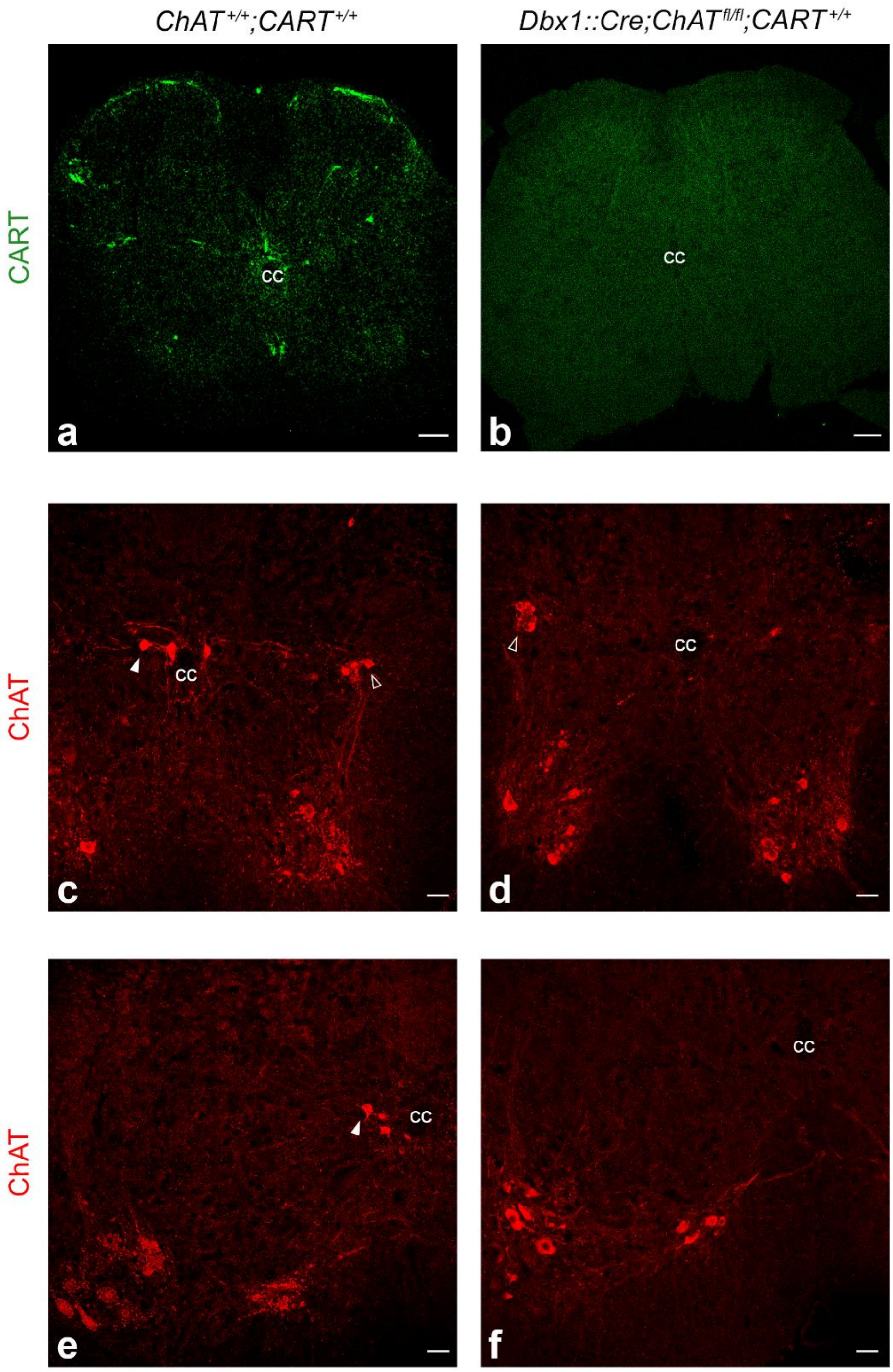


**Supplementary Fig. 5: Confirmation of the absence of CART peptide in *Cartpt* knockout mice and efficient removal of ChAT from V0c neurons**

(**a**) Immunohistochemistry for the CART peptide in upper lumbar spinal cord cross section in wild type (*$CART^{+/+}$;$ChAT^{+/+}$*) mice, reveals strong signal. (**b**) Immunofluorescence for the CART peptide in upper lumbar spinal cord cross section of *Dbx1::Cre;$ChAT^{fl/fl}$;$CART^{KO/KO}$* mice, reveals the complete absence of signal, confirming the total KO of the *Cartpt* gene. (**c**) Immunohistrochemistry for ChAT in thoracic spinal cord cross section of wild type (*$CART^{+/+}$;$ChAT^{+/+}$*) mice, reveals ChAT+ Dbx1 derived V0c interneurons (white solid arrowhead) as well as ChAT+ SPNs (empty arrowhead). (**d**) In *Dbx1::Cre;$ChAT^{fl/fl}$;$CART^{KO/KO}$* mice the Dbx1 derived V0c cholinergic interneurons are no longer ChAT+ while SPNs are still ChAT+ (empty arrowhead), proving the successful conditional KO of the *ChAT* gene specifically in V0c interneurons. (**e**) In mid lumbar sections of wild type (*$CART^{+/+}$;$ChAT^{+/+}$*) mice SPNs are not present, therefore only ChAT+ Dbx1 derived V0c interneurons are detected (white solid arrowhead). (**f**) In In *Dbx1::Cre;$ChAT^{fl/fl}$;$CART^{KO/KO}$* mice a complete absence of ChAT+ signal in the cc region reveals the successful conditional KO of the *ChAT* gene from V0c interneurons. cc: central canal. Scale bars: 50 µm.

**Supplementary Table 1**. Effects of CART or Muscarine on the intrinsic properties of fast motor neurons. The number of motor neurons studied, and animals used for each condition are included in parentheses in row 1 (MNs, animals). Data are presented as mean±SD (min, max). Statistics: data were analyzed using a repeated-measures ANOVA. F and p values are reported for each statistical test. Asterisks denote significance from Holm-Sidak post hoc comparisons, * p<0.05, ** p<0.01. *** p<0.001, **** p<0.0001. Red text signifies a significant decrease and green text a significant increase.

| **Fast Motor Neurons** | | | |
|---|---|---|---|
| **Parameter** | | **1 uM CART ($n_{MN}$=21, $n_{mice}$=11)** | **10uM Muscarine ($n_{MN}$ =12, $n_{mice}$=6)** |
| **Resting membrane potential (mV)** | | | |
| | Baseline | -64.1±4.1 (-71,-58) | -63.2±4.0(-67,-48) |
| | Drug | -66.6±4.1(-72,-59)**** | -59.3±6.1(-67,-48)** |
| | Wash | -66.4±4.3(-72,-60)** | -64.8±3.2(-68,-58) |
| | ANOVA (F,p) | 19.6, 0.00002 | 14.0,0.0005 |
| **Input resistance (MΩ)** | | | |
| | Baseline | 25.3±10.5(8.2,40.1) | 23.0±10.8(8.5,45.3) |
| | Drug | 27.6±10.1(13.4,50.0)* | 23.8±11.3(7.5,42.5) |
| | Wash | 29.7±14.2(11.6,62.2)** | 24.0±9.0(12.4,35.5) |
| | ANOVA (F,p) | 7.6, 0.003 | 6.1,0.02 |
| **Recruitment current on ramp (pA)** | | | |
| | Baseline | 534±182(295,933) | 491±233(205,868) |
| | Drug | 452±144(175,727)* | 493±286(170,1216) |
| | Wash | 515±201(175,911) | 660±399(212,1290) |
| | ANOVA (F,p) | 3.5,0.05 | 3.9,0.12 |
| **First spike threshold on ramp (mV)** | | | |
| | Baseline | -38.4±3.3(-44.7,-31.5) | -39.5±2.4(-42.9,-34.9) |
| | Drug | -38.9±4.2(-44.9,-32.2) | -41.3±2.7(-46.7,-38.2) |
| | Wash | -38.3±3.9(-45.4,-33.0) | -39.8,2.9(-44.6,-36.3) |
| | ANOVA (F,p) | 0.3,0.7 | 5.4,0.08 |
| **Action potential rise time (ms)** | | | |
| | Baseline | 0.61±0.08(0.47,0.79) | 0.56±0.14(0.2,0.75) |
| | Drug | 0.64±0.03(0.47,0.99) | 0.62±0.1(0.43,0.83) |
| | Wash | 0.65±0.1(0.48,0.80) | 0.62±0.1(0.45,0.80) |
| | ANOVA (F,p) | 1.7,0.2 | 2.9,0.13 |
| **Action potential half width (ms)** | | | |
| | Baseline | 0.69±0.15(0.47,0.98) | 0.60±0.1(0.46,0.76) |
| | Drug | 0.70±0.14(0.49,0.93) | 0.65±0.1(0.47,0.78) |
| | Wash | 0.71±0.16(0.49,0.96) | 0.62±0.06(0.54,0.69) |
| | ANOVA (F,p) | 3.2,0.08 | 5.2,0.02 |
| **Afterhyperpolarization amplitude (mV)** | | | |
| | Baseline | 6.7±1.6(3.7,10.1,) | 6.8±2.1(3.9,9.4) |
| | Drug | 7.7±2.2(4.3,12.1,)** | 3.5±2.0(1.1,8.1)**** |
| | Wash | 8.2±2.4(4.0,12.9)*** | 6.2±2.9(1.7,10.1) |
| | ANOVA (F,p) | 12.1, 0.001 | 19.7,0.0002 |
| **Afterhyperpolarization half width (ms)** | | | |
| | Baseline | 46.9±13.6(29.8,76.8) | 37.2±12.3(20.6,59.9) |
| | Drug | 49.3±14.9(33.4,82.2)* | 31.0±3.5(9.8,52.3) |
| | Wash | 49.4±14.9(32.0,83.5) | 33.2±13.8(16.8,61.3) |
| | ANOVA (F,p) | 5.3,0.02 | 5.1,0.07 |
| **Minimum firing rate (Hz)** | | | |
| | Baseline | 7.7±4.1(2.1,20.1) | 8.2±3.4(4.6,14.1) |
| | Drug | 9.9±4.2(4.6,21.5)** | 10.3±5.9(3.5,26.5) |
| | Wash | 8.1±3.5(1.9,14.7) | 8.6±5.9(3.5,18.6) |
| | ANOVA (F,p) | 6.6,0.004 | 1.8,0.2 |
| **Firing rate at 2x recruitment current (Hz)** | | | |
| | Baseline | 27.8±7.5(20.6,51.8) | 29.9±10.6(16.2,55.9) |
| | Drug | 29.7±8.8(19.5,55.6)* | 42.9±13.3(26.0,66.7)** |
| | Wash | 29.5±6.7(20.6,45.4) | 38.3±16.9(10.1,64.5) |
| | ANOVA (F,p) | 3.4,0.03 | 9.2,0.006 |
| **Maximum Firing Rate (Hz)** | | | |
| | Baseline | 43.5±9.7(25.6,61.3) | 48.9±13.0(34.0,83.3) |
| | Drug | 44.3±9.9(23.3,62.5) | 58.6±16.0(38.2,95.2)* |
| | Wash | 42.9±8.5(25.3,55.6) | 56.6±14.4(37.2,84.8) |
| | ANOVA (F,p) | 1.2,0.3 | 7.6,0.03 |

**Supplementary Table 2.** Effects of CART or Muscarine on the intrinsic properties of slow motor neurons. The number of motor neurons studied, and animals used for each condition are included in parentheses in row 1 (MNs, animals). Data are presented as mean±SD (min, max). Statistics: data were analyzed using a repeated-measures ANOVA. F and p values are reported for each statistical test. Asterisks denote significance from Holm-Sidak post hoc comparisons, * p<0.05, ** p<0.01. *** p<0.001, **** p<0.0001. Red text signifies a significant decrease and green text a significant increase.

| **Slow Motor Neurons** | | | |
|---|---|---|---|
| **Parameter** | | **1 uM CART ($n_{MN}$=15, $n_{mice}$=6)** | **10 uM Muscarine ($n_{MN}$=7, $n_{mice}$=4)** |
| **Resting membrane potential (mV)** | | | |
| | Baseline | -58.6±5.7 (-72,-52) | -58.6±7.2 (-67,-48) |
| | Drug | -59.6±6.6 (-72,-46) | -57.3±6.8 (-67,-48) |
| | Wash | -60.4±5.6 (-72,-52) | -61.4±7.9 (-68,-48) |
| | ANOVA (F,p) | 0.9, 0.4 | 5.1, 0.1 |
| **Input resistance (MΩ)** | | | |
| | Baseline | 230±221 (23,781) | 54±39 (25,129) |
| | Drug | 273±21 (30,671) | 50±41 (21,141) |
| | Wash | 246±242 (28,710) | 63±56 (22,160) |
| | ANOVA (F,p) | 1.0, 0.07 | 2.0, 0.2 |
| **Recruitment current on ramp (pA)** | | | |
| | Baseline | 64.7±71.6 (20.8,326.0) | 197±190 (55,525) |
| | Drug | 60.9±61.7 (15.5,283.0) | 238±232 (86,676) |
| | Wash | 66.0±62.2 (17.7,263.0) | 243±265 (94,714) |
| | ANOVA (F,p) | 0.6, 0.4 | 3.1, 0.2 |
| **First spike threshold on ramp (mV)** | | | |
| | Baseline | -39.5±5.6(-48.5,-30.5) | -40.2±1.9 (-43.2,-37.6) |
| | Drug | -38.9±6.2(-49.0,-27.1) | -38.8±2.5 (-41.7,-34.5) |
| | Wash | -39.8±5.6(-48.3,-30.2) | -40.5±2.7 (-44.1,-37.2) |
| | ANOVA (F,p) | 0.3,0.7 | 6.5,0.2 |
| **Action potential rise time (ms)** | | | |
| | Baseline | 0.79±0.14 (0.50,1.02) | 0.67±0.18 (0.50,0.92) |
| | Drug | 0.80±0.15 (0.44,1.06) | 0.65±0.14 (0.47,0.94) |
| | Wash | 0.84±0.16 (0.64,1.14) | 0.68±0.12 (0.51,0.80) |
| | ANOVA (F,p) | 0.7, 0.5 | 0.4, 0.6 |
| **Action potential half width (ms)** | | | |
| | Baseline | 0.91±0.2 (0.50,1.16) | 0.71±0.12 (0.53,0.86) |
| | Drug | 0.94±0.2 (0.59,1.26) | 0.68±0.16 (0.47,0.93) |
| | Wash | 0.98±0.2 (0.71,1.42) | 0.66±0.14 (0.48,0.86) |
| | ANOVA (F,p) | 2.7, 0.13 | 0.6, 0.5 |
| **Afterhyperpolarization amplitude (mV)** | | | |
| | Baseline | 8.3±2.5 (3.9,11.8) | 7.1±2.7 (3.4,11.6) |
| | Drug | 8.5±3.4 (3.8,13.9) | 4.2±2.2 (1.3,7.8) |
| | Wash | 9.5±3.1 (5.4,14.8) | 5.3±3.5 (1.1,9.0) |
| | ANOVA (F,p) | 0.7, 0.5 | 4.8, 0.08 |
| **Afterhyperpolarization half width (ms)** | | | |
| | Baseline | 81.7±26.6 (50.1,151.5) | 69.5±27.3 (37.6,109.6) |
| | Drug | 84.3±27.5 (49.1(143.6) | 42.1±15.8 (13.3,61.9) |
| | Wash | 89.3±31.5 (54.3,156.8) | 44.5±25.0 (11.6,79.9) |
| | ANOVA (F,p) | 2.4, 0.07 | 4.9, 0.05 |
| **Minimum firing rate (Hz)** | | | |
| | Baseline | 6.4±1.3 (4.6,8.3) | 4.2±1.7 (1.5,6.6) |
| | Drug | 5.6±1.5 (2.4,8.4) | 5.5±1.9 (3.3,8.8) |
| | Wash | 5.9±1.3 (4.1,7.6) | 5.7±2.2 (4.0,8.9) |
| | ANOVA (F,p) | 1.8, 0.2 | 4.5, 0.2 |
| **Firing rate at 2x recruitment current (Hz)** | | | |
| | Baseline | 19.6±6.6 (8.0,30.8) | 21.1±11.3 (9.8,40.8) |
| | Drug | 19.8±7.4 (8.5,14.4) | 24.6±14.2 (12.0,48.8) |
| | Wash | 20.6±7.5 (8.7,30.6) | 21.4±15.5 (11.1,44.3) |
| | ANOVA (F,p) | 1.8, 0.09 | 1.2, 0.3 |
| **Maximum Firing Rate (Hz)** | | | |
| | Baseline | 33.5±9.0 (20.9,56.0) | 48.2±6.9 (40.7,60.2) |
| | Drug | 31.8±9.4 (16.0,52.4) | 51.3±9.8 (37.2,60.6) |
| | Wash | 33.3±8.2 (17.8,50.8) | 48.0±9.3 (36.2,58.8) |
| | ANOVA (F,p) | 0.5, 0.5 | 0.8, 0.4 |

**Supplementary Table 3.** Comparison of the number of vAChT+ terminals, M2 receptor clusters and instances of alignment between control and *Dbx1::Cre; ChAT$^{fl/fl}$; CART$^{KO/KO}$* mice.

| *vAChT + terminals number evaluation* | | | | | | | | | | | |
|---|---|---|---|---|---|---|---|---|---|---|---|
| **mice (N)** | | **motor neurons (n)** | | **MWU** | **P value** | **median** | | **mean** | | **SE mean** | |
| ctrl | exp | ctrl | exp | | | ctrl | exp | ctrl | exp | ctrl | exp |
| 3 | 4 | 61 | 64 | 1867 | 0.6730 | 6.000 | 6.000 | 6.891 | 6.639 | 0.2929 | 0.3022 |
| *M2 receptor clusters number evaluation* | | | | | | | | | | | |
| ctrl | exp | ctrl | exp | | | ctrl | exp | ctrl | exp | ctrl | exp |
| 3 | 4 | 45 | 49 | 1001 | 0.4402 | 7.000 | 7.000 | 7.178 | 6.775 | 0.3586 | 0.3395 |
| *vAChT-M2 alignment instances number evaluation* | | | | | | | | | | | |
| ctrl | exp | ctrl | exp | | | ctrl | exp | ctrl | exp | ctrl | exp |
| 3 | 4 | 45 | 49 | 963 | 0.2880 | 7.00 | 6.000 | 7.022 | 6.490 | 0.3644 | 0.3415 |

**Supplementary Table 4.** Comparison of the size of of vAChT+ terminals and M2 receptor clusters between control and *Dbx1::Cre; ChAT$^{fl/fl}$; CART$^{KO/KO}$ mice*.

| *vAChT + terminals size evaluation* | | | | | | | | | | | |
|---|---|---|---|---|---|---|---|---|---|---|---|
| **mice (N)** | | **synapses (n)** | | **MWU** | **P value** | **median** | | **mean** | | **SE mean** | |
| ctrl | exp | ctrl | exp | | | ctrl | exp | ctrl | exp | ctrl | exp |
| 3 | 3 | 250 | 228 | 25733 | 0.0666 | 2.060 | 1.950 | 2.104 | 2.010 | 0.04124 | 0.04322 |
| *M2 receptor clusters size evaluation* | | | | | | | | | | | |
| ctrl | exp | ctrl | exp | | | ctrl | exp | ctrl | exp | ctrl | exp |
| 3 | 3 | 225 | 254 | 25658 | 0.0537 | 2.030 | 2.160 | 2.110 | 2.217 | 0.04238 | 0.03988 |
